# The Pseudogene RPS27AP5 Reveals Novel Ubiquitin and Ribosomal Protein Variants Involved in Specialised Ribosomal Functions

**DOI:** 10.1101/2024.02.05.578897

**Authors:** Anna Meller, Dominique Lévesque, Jennifer Raisch, Etienne Fafard-Couture, Michelle Scott, Xavier Roucou, Francois-Michel Boisvert

**Affiliations:** Department of Immunology and Cell biology, Université de Sherbrooke, 3201, Jean-Mignault, Sherbrooke, Québec, J1E 4K8, Canada; Department of Biochemistry and Functional Genomics, Université de Sherbrooke, 3201, Jean-Mignault, Sherbrooke, Québec, J1E 4K8, Canada

**Keywords:** pseudogene/ ribosome/ translation/ ubiquitin

## Abstract

Pseudogenes, traditionally considered non-functional gene copies resulting from evolutionary mutations, have garnered attention due to recent transcriptomics and proteomics revealing their unexpected expressions and consequential cellular functions. Ubiquitin, transcribed from UBA52 and RPS27A genes, fused to ribosomal proteins eL40 and eS31, and polyubiquitin precursors encoded by UBB and UBC genes, has additional pseudogenes labeled as non-functional. However, recent evidence challenges this notion, demonstrating that these pseudogenes produce ubiquitin variants with minimal differences from the canonical sequence, suggesting a new regulatory dimension in ubiquitin-mediated cellular processes. To systematically catalogue possible Ubiquitin (Ub) and Ubiquitin-like (Ubl) variants from pseudogenes, expression data was compiled, identifying potential functional variants. Among these pseudogenes, RPS27AP5 expresses both Ubiquitin variant (UbP5) and ribosomal protein variant (S27aP5), with precursor proteins maturing through cleavage and exhibiting behavior similar to their counterparts post-translation. Notably, S27aP5 integrates into translating ribosomes, increasing the 80S monosomal ribosomal fraction and indirectly influencing p16INK4A transcriptional activation. The discovery of a functional S27a pseudogene supports the concept that a subset of ribosomes may incorporate diverse subunits for specific translational functions.

## Introduction

Pseudogenes are by definition non-functional copies of genes (1) that can further be grouped as unprocessed, processed or unitary (2,3). Unprocessed pseudogenes originate from duplication through evolution and retain their intron-exon structure (4). Conversely, processed pseudogenes emerged from retrotransposition (5), wherein a reverse-transcribed mRNA (complete or partial) integrates into a genomic location distinct from its gene of origin (4,6,7). Within the human genome, processed pseudogenes are the most abundant type of pseudogenes due to a burst of retrotransposition events in ancestral primates (8–10). Given their mRNA origin, highly transcribed and short genes are more susceptible to accumulating agreater number of pseudogenes (11,12). Processed pseudogenes, flanked by direct repeats, resemble a reverse-transcribed mRNA, lacking intronic sequences and containing a 3’ poly A sequence in their flanking regulatory elements. Pseudogenes harbour mutations (point mutation, insertion, deletion), leading to distinct amino acid sequences, frameshifts or premature stop codons compared to the original gene (13,14). Despite their historical classification as ‘junk’ DNA, recent studies have revealed the potential functionality of pseudogenes in various cellular processes (15,16). This functionality may arise from RNA– level regulation, interacting with the parental gene transcript itself (17–20), serving as sponges of miRNAs and other regulators of the parental transcript (21,22) or after translation where they function as different variants of the parental protein (23–25).

Ubiquitin (Ub) is transcribed from four different genes: *UBB*, *UBC, UBA52* and *RPS27A.* The first two contain respectively 3 and 9 tandem repeats of Ub, while *UBA52* and *RPS27A* are encoded as fusion proteins with one Ub and L40 (eL40) or S27a (eS31) ribosomal proteins, respectively (26,27). By attaching Ub to substrate proteins, they play a role in regulating various cellular processes, including proteasomal degradation (28–32). Ubiquitin-like proteins (Ubls) represent a diverse class of small proteins that share structural similarities with ubiquitin, a well-known cellular regulator involved in various cellular processes. Unlike ubiquitin, Ubls do not typically tag proteins for degradation but instead participate in a myriad of cellular functions, including protein modification, cellular signalling, and the regulation of various biological pathways (33). The ubiquitin-like protein family includes members such as SUMO (Small Ubiquitin-like Modifier), NEDD8 (Neural Precursor Cell Expressed Developmentally Down-Regulated 8), ISG15 (Interferon-Stimulated Gene 15), and Atg8 (Autophagy-related protein 8), Atg12 (Autophagy related 12), URM1 (Ubiquitin-related modifier-1), UFM1 (Ubiquitin-fold modifier 1), and FAT10 (Human Leukocyte Antigen (HLA)-F adjacent transcript 10) (34,35).

Ribosomal encoding genes constitute the largest group of processed pseudogenes, accounting for over 20% of all pseudogenes (9,36). This high prevalence raises the possibility that some of these pseudogenes are expressed and may serve functional roles(37). Ribosomal proteins, numbering 80 in eukaryotes, play a pivotal role in the translational machinery, forming ribosomes alongside ribosomal RNAs (rRNA) (38). These proteins assemble into the two major ribosomal components, the small (40S) and the large (60S) subunits, which come together to perform protein translation (38). The maturation process of the two ribosomal subunits begins with the assembly of core proteins in the nucleolus, forming a pre-ribosomal complex, followed by cytoplasmic export and final maturation (39,40). Orchestrated by a multitude of enzymes, this complex process ensures the error-free production of mature ribosomes for translation (41–44). Translating ribosomes exist either as monosomes (80S) or as polysomes, where two or more ribosomes are loaded onto the same mRNA. While 80S monosomes primarily play a role in translation initiation, they also play a role in translating key regulatory proteins and contribute to the translation of synaptic mRNAs (45,46).

Specialized ribosomes introduce an additional layer of complexity to the intricate landscape of protein synthesis. Unlike conventional ribosomes that engage in the general translation of mRNA, specialized ribosomes are tailored to participate in specific cellular processes or respond to distinct environmental cues (47–49). These ribosomal variants, often characterized by unique protein or RNA components, contribute to the regulation of gene expression and the fine-tuning of cellular functions (50). Although still in debate, the existence of specialized ribosomes broadens our understanding of the translational machinery, highlighting its adaptability to diverse cellular contexts such as development, homeostasis, and response to cellular stress (50–53).

This study aimed to comprehensively collect all recorded Ub pseudogenes to identify potentially functional variants. Our investigation revealed that the *RPS27A Pseudogene 5* (*RPS27AP5)* not only encodes a Ub variant (UbP5), but also a ribosomal variant protein (S27aP5) derived from the S27a (eS31) ribosomal protein. We explored the interactome of both pseudogene-derived proteins to elucidate their potential functions. Interestingly, S27aP5 was found to incorporate into translating ribosomes. Employing a polysome profiling assay, we observed an unexpected increase in the 80S ribosomal fraction unpon expression of this pseudogene. Further characterization through translating ribosome affinity purification (TRAP) assay, followed by RNA sequencing, unveiled that S27aP5 preferentially participate in the translation of senescence and cell proliferation-related proteins such as KDM6B and BMI1, influencing the transcription of p16^INK4A^. Notably, the expression of S27aP5 was also linked to propagation H3K27me3 at the promoter region of *CDKN2A* (p16^INK4A^), leading to a negative impact on p16^INK4A^ protein expression levels through transcriptional repression. These findings indicate that S27aP5 could play an important role in defining a subset of ribosomes with specialised function.

## Results

### Collection of Ub and Ubl pseudogenes

To identify functional Ub-containing pseudogenes, a comprehensive search was conducted in human genome databases containing both Ub and Ub-like (Ubl) pseudogenes. Based on parental gene filtration, a compiled list was generated from these pseudogene databases. Proceeding from this dataset, further curation was performed considering sequence similarity to parental gene, presence of RNA or protein level of evidence. A total of 57 Ub and 51 Ubl pseudogenes were identified(36,54,55). Some of these pseudogenes showed evidence of expression at the RNA level based on RNA-seq data, as indicated in Supplementary Tables 1-2 (25,56,57). After compiling all identified pseudogenes, potential functional candidates were selected by considering protein sequence similarities to parental genes and the presence of a Gly-Gly motif at the C-terminal, facilitating substrate conjugation(Suppl Fig 1-4)(58,59). Given the interest in identifying novel variants that can be attached to target proteins, candidates were further assessed for evidence of translation through ribosome profiling experiments. Also, since both *UBA52* and *RPS27A* parental genes encode for a Ub and a ribosomal protein, it was particularly interesting to see if any of their pseudogenes could give rise to a variant ribosomal protein as well. Despite the presence of 33 *RPS27A* pseudogenes and 16 *UBA52* pseudogenes, only 6 and 1, respectively, had sequences extending into the ribosome coding region (Suppl Table 1-2). The selection process was refined by confirming translation through ribosome profiling experiments and verifying protein-level expression using large-scale mass spectrometry datasets re-analyzed with OpenProt (56,60) and PepQuery (61,62) (Suppl Fig 5). Five potentially intriguing Ub pseudogenes: *UBA Pseudogene 6* (*UBA52P6*), *UBB Pseudogene 3* (*UBBP3*), *RPS27A Pseudogene 1* (*RPS27AP1*), *RPS27A Pseudogene 5* (*RPS27AP5*), and *RPS27A Pseudogene 11* (*RPS27AP11*), were selected for functional analysis based on these criteria (Fig.1a-b). Each pseudogene exhibited 8, 14, 8, 3, and 6 amino acid differences, respectively, within the canonical Ub coding region. Remarkably, *RPS27AP5* not only encoded a Ub variant (UbP5) but also a full-length ribosomal protein S27a variant (S27aP5) within the same open reading frame, mirroring the parental *RPS27A* gene.

To assess the potential conjugation of these Ub variant proteins to substrates, HeLa cells were transfected with HA-tagged pseudogene-derived Ub constructs. Immunoblot analysis was conducted to determine the expression and conjugation of the respective Ub pseudogenes (Figure 1c). For comparison, Ub and the previously identified functional pseudogene Ub^KEKS^ (25) were included in the assay to evaluate modification patterns and localization. The expression levels of pseudogene proteins varied significantly, sometimes approaching the limit of detection (Fig.1c). To investigate the stability of the substrate proteins and their potential targeting for proteasomal degradation, transfected cells were treated with the proteasomal inhibitor MG132 (Fig.1c). HA immunoblots revealed bands for UbP5 and RPS27aP11 at different molecular weights above the size of monomeric Ub, indicating successful conjugation to substrate proteins (Fig.1c). Higher molecular weight protein accumulation was observed with these two candidates after MG132 treatment, suggesting involvement in substrate targeting for degradation.

**Figure 1.**
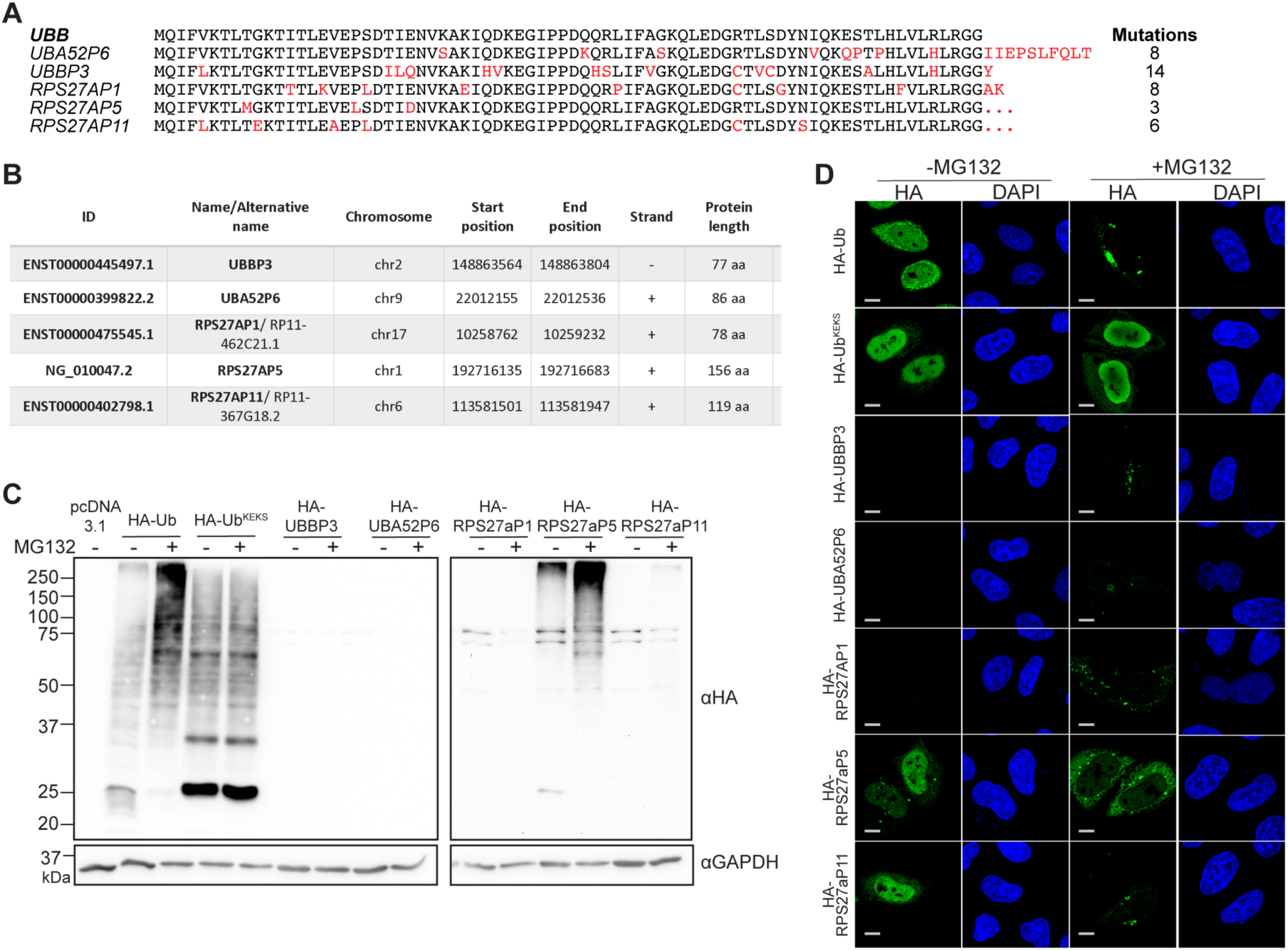
Identification of potentially functional Ub pseudogenes. A. Amino acid sequence alignment of proteins coded by selected pseudogenes. The red colour indicates differences compared to the Ub sequence. B. ID, chromosomal localization and protein length of selected pseudogenes with Gly-Gly motifs, RNAseq, ribosome profiling and mass-spectrometry evidence (assembly version: GRCh38). C. Immunoblot analysis of HA tagged Ub, Ub^KEKS^ and selected pseudogenes-encoded Ub variants, both with or without treatment with 20 µM MG132 (N=3). Total cell lysates were separated by SDS-PAGE, and proteins were detected with the HA antibody. D. Immunofluorescence analysis of HA tagged Ub, Ub^KEKS^, and selected pseudogenes-encoded Ub variants with or without treatment with 20 µM MG132 (N=3). Proteins were detected with the HA antibody and the nuclei were labelled with DAPI. Scale bars represent 10 μM.

Immunofluorescence analysis was also performed to determine protein localization under normal conditions and following MG132 treatment (Fig.1d). While UbP5 and RPS27aP11 primarily showed nuclear localization similar to Ub and Ub^KEKS^ under normal conditions, UBBP3, UBA52P6, and RPS27aP1 were undetectable. Only a faint signal, indicating accumulation in granules, was observed in response to MG132. Notably, UBA52P6 and UBBP3, despite having protein-level evidence, were challenging to detect after overexpression. In case of UBA52P6, potentially due to tissue specificity, exhibiting the highest expression levels in testes (Suppl Fig 2b). Among all investigated pseudogene-derived proteins, only UbP5 and RPS27aP11 were expressed abundantly and demonstrated the ability to be conjugated to substrates. They displayed a similar capacity to target proteins for degradation as wild-type Ub. Due to its robust overexpression, UbP5, and the potential expression of a ribosomal protein (S27aP5) from RPS27AP5, further experiments were focused on these gene products.

### Characterization of RPS27AP5, a pseudogene encoding both a Ub and a ribosomal variant

To validate the results obtained from the database search, both the genomic and mRNA sequences of *RPS27AP5* were confirmed through PCR amplification and subsequent sequencing of genomic DNA (gDNA) and complementary DNA (cDNA). The S27aP5 protein closely resembles S27a, with only four differing amino acids: K89R, V102L, R116C, and R118H (Fig.2a). Unlike *RPS27A*, *RPS27AP5* lacks intronic sequences and possesses a poly-A tail, indicating its status as a processed pseudogene resulting from the insertion of reverse-transcribed mRNA into the genome (6). The presence of *RPS27AP5* mRNA was confirmed through reverse transcription and gene-specific PCR in at least three different cell lines (HeLa, U2-OS, and HEK293a) (Suppl Fig.6).

To investigate whether S27aP5 is translated from the *RPS27AP5* mRNA and undergoes maturation similar to S27A through deubiquitylating enzymes(63), immunoblot and immunofluorescence analyses were conducted. Both proteins were transiently expressed in cells, each carrying an N-terminal HA tag and a C-terminal Myc tag for the detection of Ub/UbP5 and S27a/S27aP5 proteins (Fig.2b). Immunofluorescence analysis demonstrated that both proteins are translated from the pseudogene sequence and localized separately, confirming cleavage after translation. (Fig.2c). The localization pattern of S27aP5 closely resembled that of wild-type S27a, where Ub is distributed throughout the entire cell, and ribosomal protein S27a is predominantly found in the nucleoli, attached to the rough endoplasmic reticulum (ER), or freely present in the cytosol(64,65). This observation suggests that the ribosomal pseudogene protein S27aP5 undergoes similar processing and transport mechanisms as S27a after translation. S27a is a component of the 40S small ribosomal subunit, composed of a total of 33 proteins(66,67). It contains C4-type zinc finger domains and is situated on the external surface of the ribosome, directly binding to the 18S ribosomal RNA (63). Within the complex, deubiquitylation of S27a at K113 by USP16 has been demonstrated to be crucial for 40S subunit maturation, implying a role for this protein in the quality control mechanism (68).

The immunoblot analysis indicated that both UbP5 and S27aP5 exhibited lower expression compared to Ub and S27a (Fig.2d). To investigate whether the observed differences are attributed to the UbP5 component of the S27aP5 protein, chimeric sequences were generated by swapping either the Ub or the ribosomal part of the pseudogene (Fig.2d). Intriguingly, substituting the UbP5 sequence with Ub did not enhance the expression level of S27aP5. Similarly, replacing the Ub sequence with UbP5 had no effect on the S27a protein level (Fig. 2c). These findings suggest that the regulation of S27aP5 expression occurs after translation and cleavage, independent of translation efficiency.

**Figure 2.**
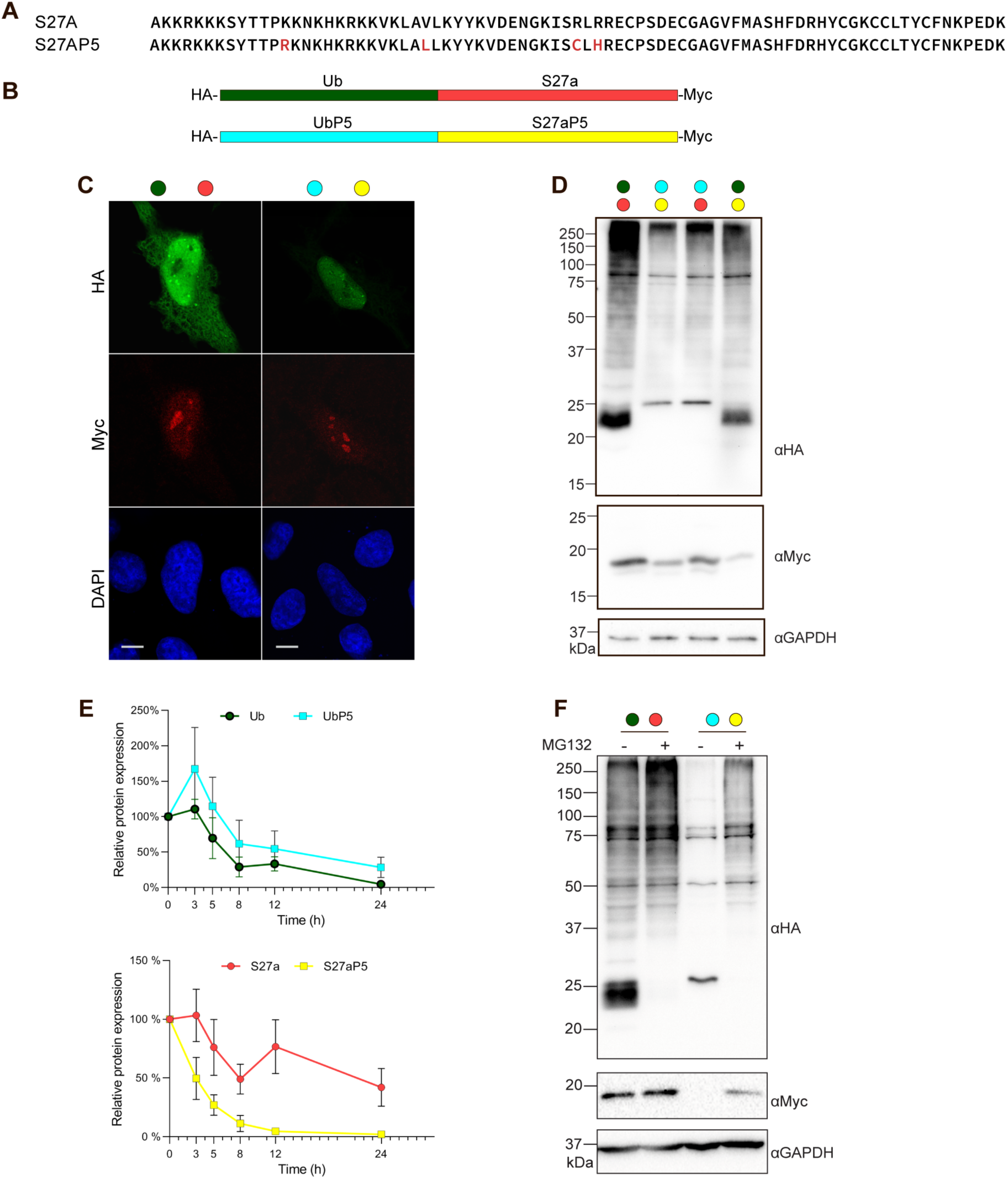
RPS27AP5 encodes for a Ub and a ribosomal (S27a) protein. A. Amino acid sequence alignment of the ribosomal protein encoded by RPS27A and RPS27AP5. Differences from the parental protein are highlighted in red. B. Schematic representation of the expression construct of RPS27a and RPA27aP5. C. Immunofluorescence analysis of RPS27a and RPA27aP5 (N=3). Proteins were detected with the HA antibody, and the nuclei were labelled with DAPI. Scale bars represent 10 μM. D. Immunoblot visualization of HA-Ub-S27A-Myc and HA-UbP5-S27AP5-Myc and chimeric constructs of HA-UbP5-S27A-Myc and HA-Ub-S27AP5-Myc. Total cell lysates were separated by SDS-PAGE, and proteins were detected with HA and Myc antibodies. E. Protein stability assay of RPS27a and RPS27aP5. Cells overexpressing either construct were treated with 15 µg/ml CHX (cycloheximide) and were harvested at the indicated timepoints (N=3). Total cell lysates were separated by SDS-PAGE and proteins were detected with the HA and Myc antibodies. Protein ratios were calculated by normalization to total protein (Ponceau) levels, with untreated samples considered as 100%. Error bars represent SEM. F. Rescue experiment for proteasomal degradation. Cells overexpressing RPS27a or RPS27aP5 were treated with 20 µM MG132 to prevent degradation (N=3). Total cell lysates were separated by SDS-PAGE and proteins were detected with the HA and Myc antibodies.

To assess whether these differences are linked to protein stability, cells expressing either HA-Ub-S27a-Myc or HA-UbP5-S27aP5-Myc constructs were treated with cycloheximide (CHX) to halt new protein synthesis and harvested at different time points. Immunoblot analysis with HA and Myc antibodies revealed that while UbP5 stability was similar to Ub, S27aP5 levels started decreasing more rapidly after 3 hours, indicating a shorter half-life compared to the parental protein (Fig. 2e). Given that this shorter half-life might be associated with proteasomal degradation, cells expressing either HA-Ub-S27a-Myc or HA-UbP5-S27aP5-Myc were treated with MG132 to inhibit degradation. Immunoblot analysis demonstrated that although the treatment increased S27aP5 protein levels, it did not fully restore them to levels comparable to S27a (Fig. 2f), suggesting that the stability of S27aP5 is partially regulated through the proteasome.

### Identification of RPS27aP5 interactors and targets

To explore the potential functions of the proteins encoded by *RPS27AP5*, affinity purification was conducted using cells overexpressing either HA-Ub-S27a-Myc or HA-UbP5-S27aP5-Myc constructs. Cell lysates were divided into three fractions, and immunoprecipitation of Ub/UbP5 or S27a/S27aP5 was performed using HA or Myc antibodies with an additional untransfected control sample for each antibodies. The pulldown was carried out under non-denaturing conditions to capture both interactors and substrates, particularly in the case of Ub and UbP5. Recovered proteins were identified through mass spectrometry, employing a DIA acquisition method. Immunoprecipitation of the Ub portion of both genes confirmed that UbP5 interacts with proteasomal proteins and several components of the Ub conjugating machinery, supporting the hypothesis of a proteasome targeting function (Fig. 3a-b). However, certain proteins such as EIF6, a translation initiation factor, and OTULIN, a deubiquitinase enzyme, were enriched in the UbP5 pulldown, while others like RNF123, ITCH, XIAP, and RBCK1 E3 Ub ligases were enriched in the Ub pulldown exclusively (Fig. 3c). This suggests that although UbP5 shares very similar functions with Ub, these two Ub variants may have subtle differences in target proteins, interacting with distinct sets of proteins. To validate the obtained results, a co-immunoprecipitation assay was conducted using OTUB1 as a common interactor for both Ub and UbP5. Cells overexpressing Flag-OTUB1 and HA-Ub or HA-UbP5 were immunoprecipitated with a Flag antibody, and the presence of Ub or UbP5 was detected by immunoblot using an HA antibody (Fig. 3d). Proteins modified by Ub or UbP5 interacting with OTUB1 were identified with similar intensities, affirming their comparable association with the common interactor.

**Figure 3.**
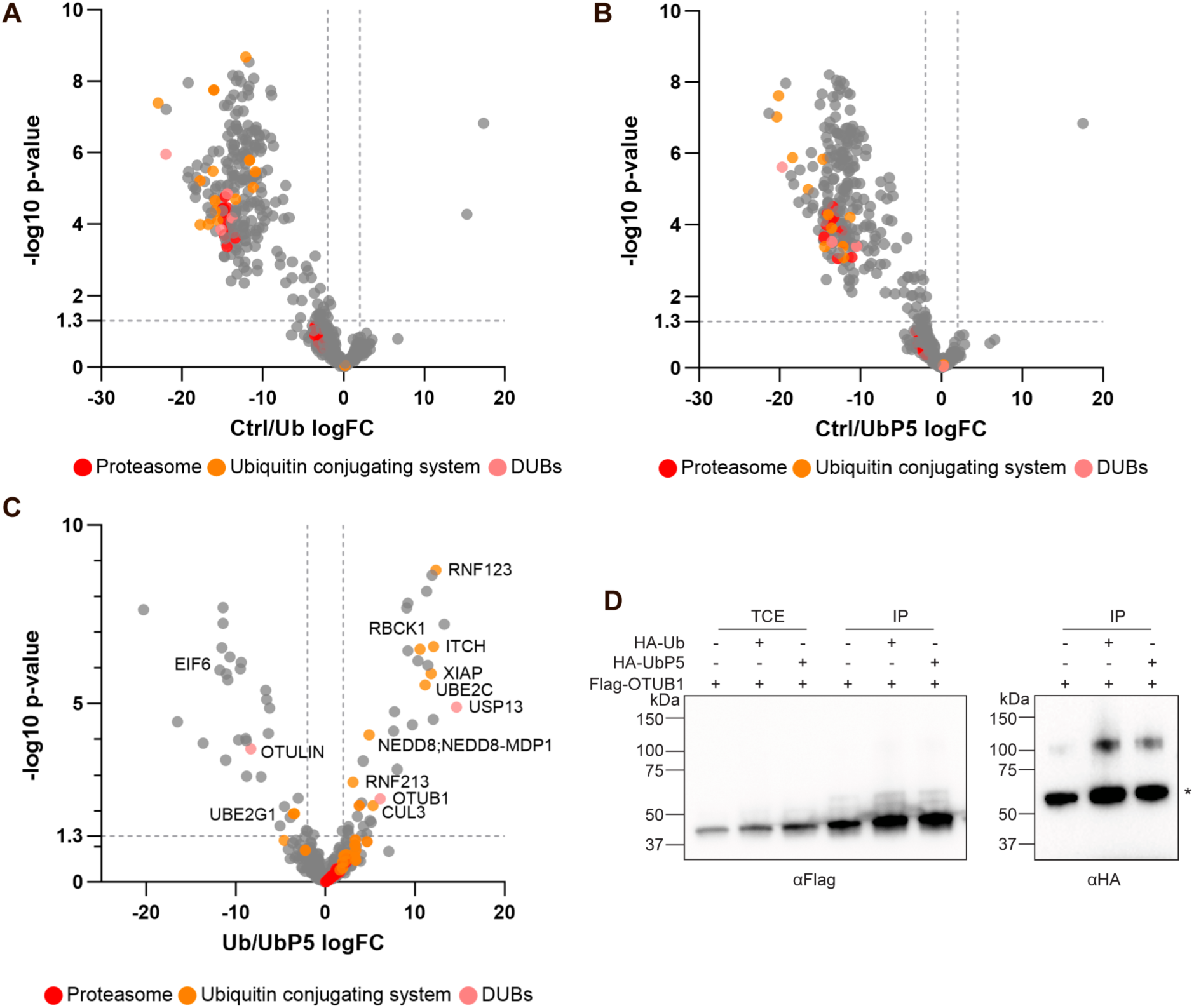
Comparison of Ub and UbP5 interactome. A-B. Comparison of proteins interacting with either Ub or UbP5 and untransfected control cells (Ctrl) (N=3). Proteins related to the proteasome are labelled in red, proteins part of the Ub conjugating system are labelled in yellow, and deubiquitylating proteins are labelled in pink. C. Comparison of proteins interacting with Ub and UbP5, enriched in immunoprecipitation performed in cells expressing HA-Ub or HA-UbP5 (N=3). Proteins related to the proteasome are labelled in red, proteins part of the Ub conjugating system are labelled in yellow, and deubiquitylating proteins are labelled in pink. D. Immunoprecipitation assay with Flag-OTUB1 as an interacting partner for HA-Ub and HA-UbP5 (N=3). Cells were co-transfected with OTUB1 and RPS27a or RPS27aP5. Total cell lysates and immunoprecipitated proteins were separated by SDS-PAGE and labelled proteins were visualised using HA and Flag antibodies. Asterix (*) indicates a non-specific band.

**Figure 4.**
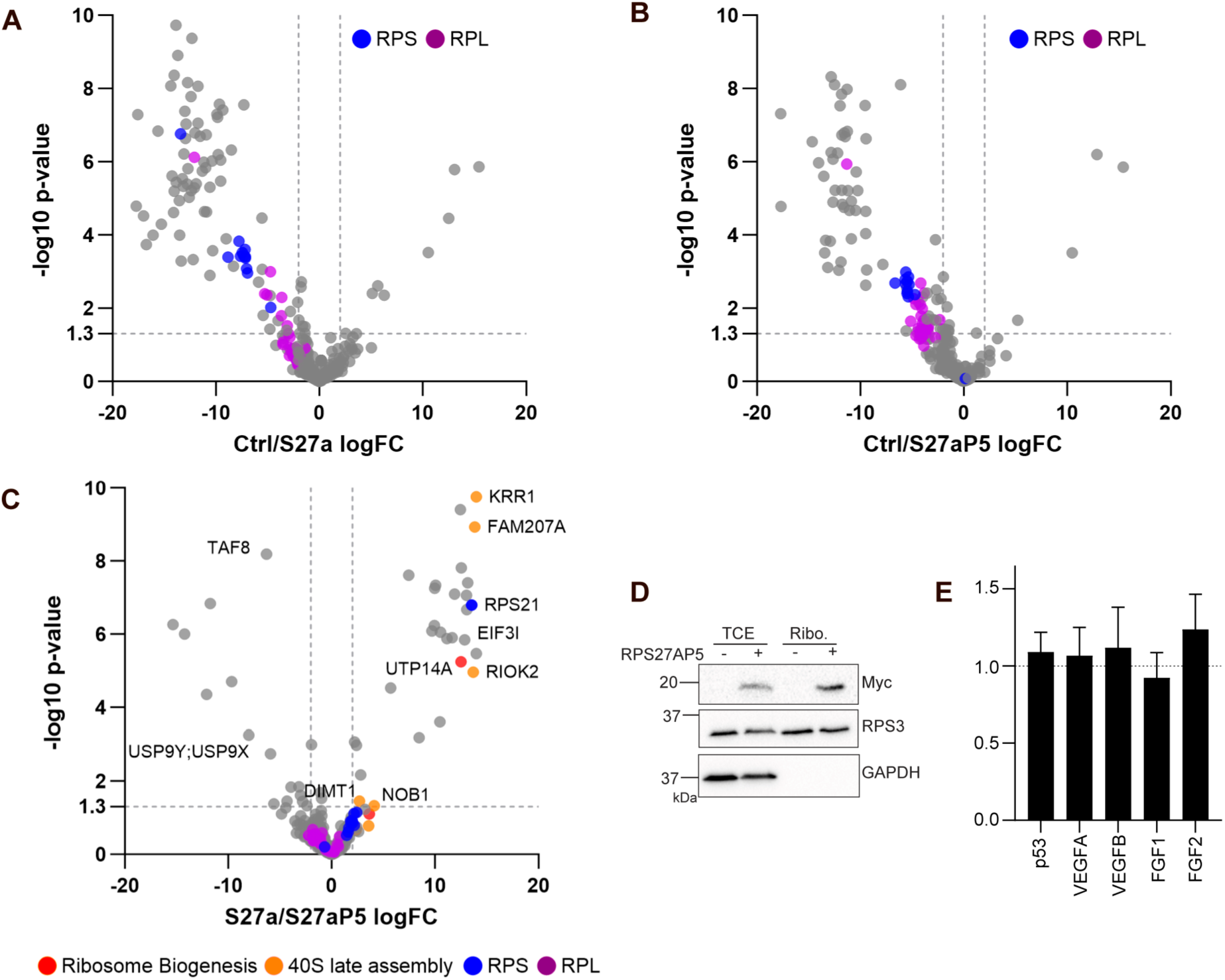
Comparison of S27a and S27aP5 interactome. A-B. Comparison of proteins interacting with either S27a or S27aP5 and untransfected control cells (Ctrl) (N=3). Proteins that are part of the large ribosomal subunit (60S) are labelled in purple, small ribosomal subunit proteins (40S) are labelled in blue. C. Comparison of proteins interacting with S27a and S27aP5, enriched by immunoprecipitation performed in cells expressing Myc-S27a or Myc-S27aP5 (N=3). Proteins that are part of the large ribosomal subunit (60S) are labelled in purple, small ribosomal subunit proteins (40S) are labelled in blue, ribosome biogenesis related proteins are labelled in red, and proteins related to late 40S assembly are labelled in yellow. D. Ribosome extraction from un-transfected and S27aP5 expressing HeLa cells. Total cell lysates, supernatant and ribosome containing pellet fractions were separated by SDS-PAGE and labelled proteins were detected using the Myc antibody. E. Luciferase assay for detection of cap-independent translation efficiency of cell expressing either RPS27a or RPS27aP5. Cells were co-transfected with RPS27a or RPS27aP5 and p53, VEGFA, VEGFB, FGF1 or FGF2 IRES dual expression plasmids with Renilla and firefly luciferases (N=3). Luminescence intensities were normalised to Renilla expression within each sample and the results were visualized as a ratio considering RPS27a as 1.

Interactome analysis of ribosomal proteins S27a and S27aP5 revealed interactions with members of both the small and large ribosomal subunits (Fig. 4a-b), suggesting that S27aP5 can be incorporated into an 80S monosome. This is further supported by the presence of interacting proteins shared between parental S27a and S27aP5 that are components of the ribosome (Fig. 4c). However, certain proteins related to ribosomal biogenesis and 40S maturation were notably absent from the S27aP5 interactome. Specifically, RIOK2, a protein involved in 40S maturation(69), as well as KRR1, UTP14A, and FAM207A, proteins involved in ribosome biogenesis (70–72), were exclusively detected in the S27a interactome. This suggests a potential alteration in the maturation process of ribosomes containing S27aP5 (Fig. 4c). Intriguingly, the deubiquitylating enzyme USP9X, implicated in ribosome stalling and collision (73), was found to be enriched only in the S27aP5 interactome.

To confirm the potential integration of S27aP5 into ribosome complexes, ribosome extraction was performed. Cells overexpressing S27aP5 were treated with cycloheximide (CHX) to stall ribosomes during translation, and protein lysates were layered over a sucrose cushion. Following centrifugation, the resulting pellet containing assembled ribosomes and ribosomal subunits was subjected to immunoblot analysis. S27aP5 was detected in the ribosome fraction (Fig. 4d), confirming its ability to integrate into a 40S subunit or a ribosome complex. Despite the observed lack of interaction with certain crucial ribosome-related factors, the investigation of Internal Ribosome Entry Site (IRES)-mediated cap-independent translation using a luciferase assay with p53, VEGFA, VEGFB, FGF1, and FGF2 IRES in cells overexpressing either S27a or S27aP5 revealed no defects in translation efficiency (Fig. 4e). These results indicate that the overexpression of S27a has no adverse effect on cap-independent translation.

### Role of S27aP5 within the ribosome

As ribosome extraction primarily relies on the pelleting of large protein complexes, it lacks the ability to distinguish between different ribosomal species such as 40S, 60S, 80S ribosomes, or polysomes. To enhance resolution and discern whether S27aP5 integrates into actively translating ribosomes, a polysome profiling assay was implemented (74,75). Cells overexpressing S27a or S27aP5 were treated with CHX, and protein lysates were subjected to density separation using a sucrose gradient, allowing the isolation of individual ribosomal subunits, monosomes, and polysomes, representing two or more ribosomes translating the same RNA molecule. The separated ribosomal fractions were collected and visualized using a gradient fractionator (Fig. 5a). Precipitated protein fractions corresponding to 40S, 60S, 80S, and polysomal components were analyzed for the presence of S27a and S27aP5 through immunoblotting with a Myc antibody (Fig. 5b).

**Figure 5.**
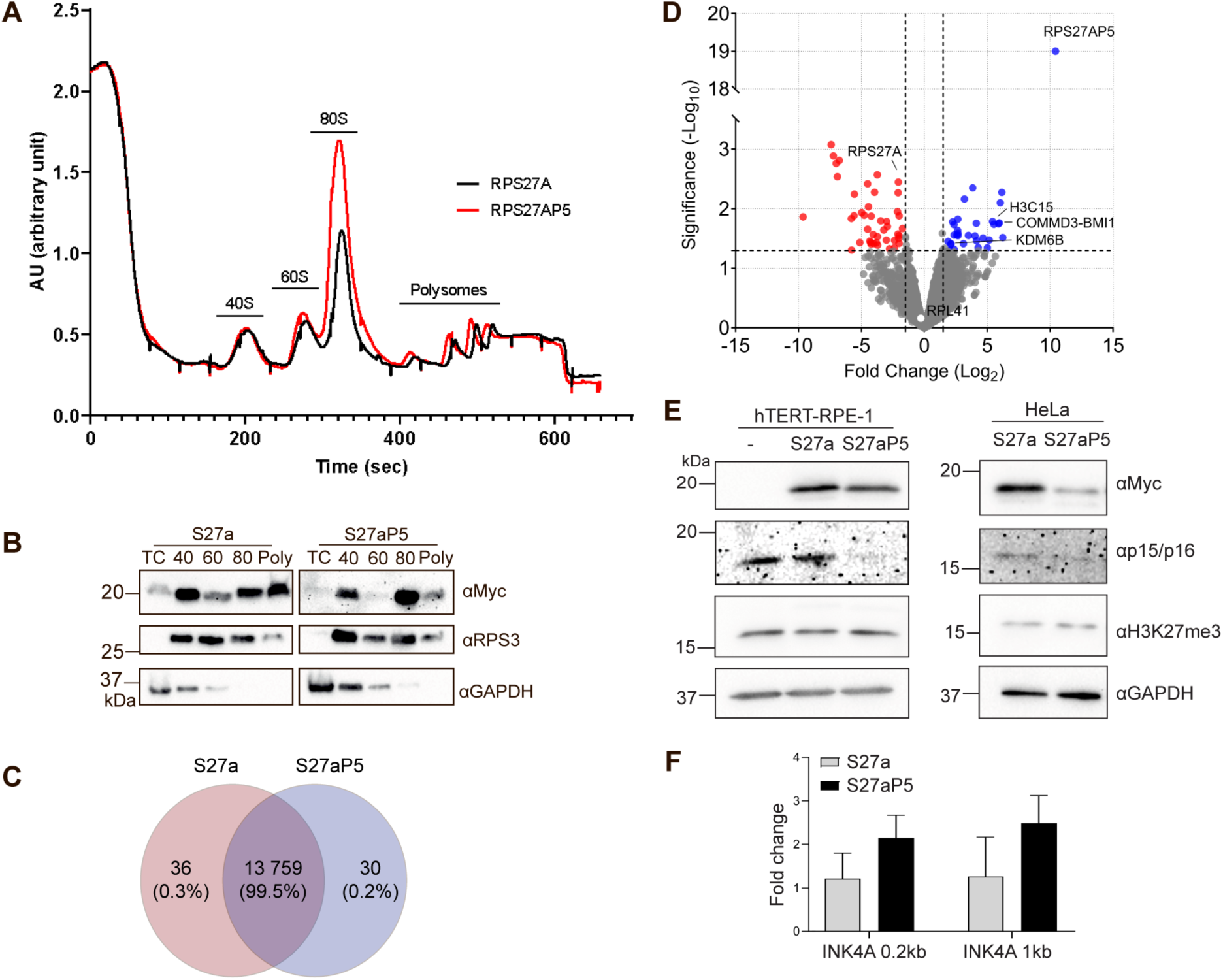
Role of S27aP5 within the ribosome. A. Polysome profiling assay with cells expressing either Myc-S27a or Myc-S27aP5 proteins (N=2). Cell lysates expressing RPS27a or RPS27aP5 and treated with CHX (cycloheximide) were layered over a continuous sucrose gradient, and proteins were separated by ultracentrifugation. Separated ribosomal units were visualised using a gradient fractionator. B. Immunoblot analysis of precipitated fractions corresponding to the 40S, 60S, 80S ribosomal and polysomal proteins collected from polysome profiling assay (N=3). Fractions from polysome profiling were collected, and proteins following precipitation were separated by SDS-PAGE. Ribosomal proteins were revealed with the Myc and RPS3 antibodies. C. Venn diagram representation of RNA-Seq results of total RNAs captured by TRAP assay using either Myc-RPS27a or Myc-RPS27aP5 as bait (N=3). Cells expressing RPS27a or RPS27aP5 and treated with CHX (cycloheximide) were subjected to Myc immunoprecipitation. Ribosome-bound RNAs were recovered and analysed using RNA-seq. Identified RNAs bound to either RPS27a or RPS27aP5 were compared using a Venn diagram. D. Comparison of RNA-Seq results of protein-coding RNAs identified from the TRAP assay using either Myc-RPS27a or Myc-RPS27aP5 as bait (N=3). Log2 Fold Change threshold=1 and Significance cut-off is at 1.3 (0.05% FDR). E. Immunoblot analysis of HeLa and hTERT-RPE-1 cell extracts. Lysates from un-transfected cells, or cells expressing either RPS27a or RPS27aP5, were separated by SDS-PAGE and detected using the Myc, H3K27me3 and p15/16 antibodies. F. qPCR quantification of H3K27me3 levels at two different sites within the promoter region of *INK4A* in wild type HeLa cells or cells expressing S27a or S27aP5 proteins. Results were normalised to a spike-in DNA control and are shown relative to the un-transfected control sample used. Error bars correspond to standard deviation (N=3).

Bands corresponding to S27aP5 were evident not only in the monosomal 80S fraction but also in the polysome fraction, confirming the integration of S27aP5 into translating ribosomes. Notably, the majority of S27aP5 was found in monosomes (Fig. 5b). Furthermore, while overexpression of the parental S27a protein did not impact the 80S fraction, a slight increase in monosomes was observed in S27aP5-overexpressing cells on the polysome profiling chromatogram (Fig. 5a). These observations may suggest a potential disruption in ribosomal function due to the substitution of the wild-type S27a protein with the pseudogene or a maturation defect resulting in the removal of S27aP5-containing 80S ribosomes from translation (76). It is also conceivable that ribosomes containing S27aP5 exhibit a preference for monosomes, participating in a specialized ribosomal function related to 80S translation(45,77).

### Identification of proteins translated by S27aP5 containing ribosomes

As the overexpression of S27aP5 impacted the 80S fraction, a Translating Ribosome Affinity Purification (TRAP) assay was employed to investigate whether ribosomes containing this variant exhibit a preference for translating a specific subset of mRNA. Cells overexpressing S27a or S27aP5 were treated with CHX, and protein extracts were subjected to immunoprecipitation using Myc nanobody-coupled beads. The captured translating ribosome-bound RNA was recovered and analyzed using RNA-Seq. Comparison of identified RNAs between the two subunits revealed that the majority of RNAs were similar in both pulldowns, with 36 and 30 RNAs interacting uniquely with S27a or S27aP5-containing ribosomes, respectively (Fig. 5c). This confirms that the functions of S27a and S27aP5 are very similar under normal conditions.

To further explore the differences in translated RNAs between ribosomes containing S27a and S27aP5, DESeq2 differential analysis was performed (Fig. 5d). Despite the expectation that S27aP5, primarily associated with monosomes, might preferentially interact with shorter mRNAs, both S27a and S27aP5 ribosomes exhibited similar associations with RPL41, a ribosomal protein with a 78-nt-long coding sequence known to accommodate only one ribosome (Fig.5d)(78). The mRNA populations exhibiting enriched association with the S27a or S27aP5 containing ribosomes has further been analysed for 5’ UTR length, number of upstream open reading frames (uORFs) and half-life (Suppl Fig 7). The analyses showed that mRNAs associating with S27aP5 had a 5’UTR length <500 bp, with their main half-life values on the slightly longer end. Due to the small number of transcripts however, there was no significant difference observed compared to mRNAs associating with S27a (Suppl Fig 7). This observation suggests that ribosomes containing S27aP5 may have a specialized translational function related to 80S monosomes.

Gene ontology analysis revealed an enrichment in three genes related to senescence (*KDM6B*, *COMMD3-BMI1*, and *H3C15*) in ribosomes containing S27aP5 (Fig. 5d). During senescence, KDM6B (JMJD3), a histone demethylase, is responsible for the removal of H3K27me2/3 (79–81) from *CDKN2A* (p16^INK4A^), leading to its transcriptional activation (82–85) and its decrease is related to the attenuation of *CDKN2A* and *CDKN1A* transcription (86). On the other hand, BMI1, along with RING1 (RING1A), RNF2 (RING1B) Ub ligases, and CBX proteins, is part of the Polycomb Repressor Complex 1 (PRC1) (87,88), acting as a transcriptional repressor of *CDKN2A*(89,90). Notably, S27a protein has previously been linked to cell cycle arrest by regulating the MDM2-p53 loop. S27a can either bind RPL11, preventing its binding to MDM2 and facilitating MDM2-mediated ubiquitylation (91), or it can bind MDM2, preventing p53 ubiquitylation (92).

To further investigate whether S27aP5 has a direct impact on p16^INK4A^ protein expression, both S27a and S27aP5 were overexpressed in hTERT RPE-1 cells (with wild-type p53 status) and in HeLa cells. Protein levels of p16^INK4A^ and H3K27me3 were assessed using immunoblot analysis (Fig. 5e). While the level of H3K27me3 remained unaffected by the overexpression of either RPS27a or RPS27aP5, the protein level of p16^INK4A^ exhibited a decrease in cells overexpressing S27aP5 (Fig. 5e). To confirm that the reduction in p16^INK4A^ levels is associated with the regulation of histone methylation by KDM6B, a CUT and RUN (Cleavage Under Targets and Release Using Nuclease) assay was performed, targeting H3K27me3 sites, followed by qPCR on the *CDKN2A* promoter (Fig. 5f). The results obtained indicated that while overexpression of RPS27a did not impact H3K27me3 levels on the investigated *CDKN2A* promoter regions, overexpression of S27aP5 slightly increased H3K27me3 on these sites. This observation suggests a potential transcriptional repression of p16^INK4A^ due to the augmented H3K27me3 levels induced by S27aP5. Consequently, S27aP5 appears to influence the expression of p16^INK4A^ through indirect transcriptional regulation.

## Discussion

In this study, an extensive collection of Ub and Ubl pseudogenes was systematically generated to discern functional proteins among them. Among the chosen ubiquitin variant candidates, only two, namely RPS27AP11 and RPS27AP5 (UbP5), exhibited detectable Ub expression levels and demonstrated conjugation to substrate proteins, suggesting the existence of additional Ub variants (Fig. 1). The modification of proteins by these Ub pseudogenes directed the protein substrates towards proteasomal degradation, as evidenced by the accumulation of higher molecular weight proteins upon MG132 treatment, indicative of polyubiquitylation (Fig. 1). This implies the potential of additional Ub variants that target proteins for proteasomal degradation, potentially in a cell type-, cell cycle stage-, or stress-dependent manner, similar to SUMO proteins(93,94). Of particular interest, RPS27AP5 encoded not only a Ub protein (UbP5), but also a ribosomal S27a protein (S27aP5) from its pseudogene, providing an avenue for investigating a potential ribosomal protein variant.

UbP5 and S27aP5 were observed to undergo maturation through post-translational cleavage and exhibited independent localization, akin to the parental *RPS27A* gene-encoded proteins (Fig. 2), suggesting the existence of two functional proteins derived from a single pseudogene. Subsequent interactome analysis indicated that UbP5 interacts with proteasomal proteins and targets similar substrates as Ub (Fig. 3). This finding raises the question of whether UbP5 is a variant without a specialized cellular function or if its specificity is linked to the ribosomal subunit protein, possibly involved in the specific modification of S27aP5. Supporting the notion of an S27a paralogue, S27aP5 was found to interact with both small and large ribosomal subunits, with integration into ribosomes confirmed through sucrose co-sedimentation experiments (Fig. 4). However, certain 40S maturation factors were found to interact with S27a but not with S27aP5. This discrepancy could be attributed to the lower expression and half-life of S27aP5 (Fig. 2) and the fact that the co-immunoprecipitation assay was not designed to detect interactions with maturation factors (Fig. 4). Notably, S27a ubiquitylation at K113 affects late 40S maturation, and S27aP5 has mutations in close proximity to this site, suggesting a potential maturation defect in ribosomes containing S27aP5.

Contrary to previous assumptions, the composition and stoichiometry of ribosomes are dynamic, with variants of the canonical machinery identified as potentially specialized forms (95). Functionally specialized ribosomes can arise from changes involving ribosomal proteins or ribosomal RNA (rRNA), impacting tissue-specific functions, stress responses, or specific cellular pathways (46–49). While overexpression of S27aP5 increased the 80S population (Fig. 5), it did not affect polysome quantities. To investigate whether ribosomes containing S27aP5 face difficulties in translation elongation or are part of a specialized ribosome, a TRAP assay capturing ribosome-bound RNAs combined with RNAseq was conducted (Fig. 5). Results showed that the majority of interacting RNAs were similar between ribosomes containing S27a or S27aP5, suggesting that S27aP5 containing ribosomes are properly functioning variants of the canonical complex.

However, in ribosomes containing S27aP5, certain mRNAs exhibited increased enrichment, including *KDM6B*, *COMMD3-BMI1*, and *H3C15*. Notably, these genes play roles in cell proliferation and senescence (Fig. 5). Both KDM6B (JMJD3) and BMI1 function as transcriptional regulators of p16INK4A (CDKN2A) in an antagonistic manner and are involved in H3K27 methylation. KDM6B (JMJD3) acts as a demethylase, removing H3K27me3 from p16^INK4A^ promoter regions, thereby activating its transcription. This activation leads to G1 cell cycle arrest through the inhibition of CDK4/6 (Cyclin-dependent kinase 4/6) and repression of the retinoblastoma (Rb) gene (82,96). On the other hand, the Polycomb repressor complex 2 (PRC2) member EZH2 (Enhancer of zeste homolog 2) is responsible for the trimethylation of H3K27, promoting the recruitment of PRC1, which contains BMI1. This complex, through RING1B, monoubiquitylates H2A at K119, leading to RNA Pol II (RNA polymerase II) elongation inhibition and subsequent p16^INK4A^ silencing, cell cycle progression, and proliferation (97,98). Immunoblot analysis of H2K27me3 and p16^INK4A^ levels in cells overexpressing either S27a or S27aP5 revealed that, while the overall level of H3K27me3 remained unchanged, p16^INK4A^ expression decreased in the presence of S27aP5, suggesting specific transcriptional regulation of the *CDKN2A* gene (Fig. 5). P16^INK4A^ inactivation was further confirmed using H3K27me3 CUT and RUN qPCR in cells overexpressing S27a and S27aP5 (Fig. 4). While the overall H3K27me3 level did not change in S27a overexpressing cells compared to the control, S27aP5 overexpression led to an increase in H3K27me3 at the *CDKN2A* promoter region.

In conclusion, the identification and analysis of Ub pseudogenes uncovered several potentially functional pseudogenes expressed at both the RNA and protein levels. The study of the ribosomal pseudogene protein S27aP5 revealed its impact on the monosome population and its indirect role in transcriptional silencing of p16^INK4A^. These findings support the hypothesis of ribosome specification and introduce the concept of gene regulation through alternative ribosomal proteins.

## Methods

### Data collection

The Ub encoding genes (*UBB, UBC, UBA52* and *RPS27A*) were used to search for related pseudogenes in human pseudogene databases, Pseudogene.org (GENCODE v.10) (54,99) and Pseudofam/PseudoPipe (55,100). *UBB* has five pseudogenes (*UBBP1-5*), *UBC* has four (*AC073325.2, AC108676.2, RP3-432I18.1* and *L596327.1*), *UBA52* has 11 named pseudogenes (*UBA52P1-9, AC99535.1* and *AC005165.2*) and five without names, and RPS27A has 22 named pseudogenes (*RPS27AP1-19, AC008072.3, AC079741.5* and *AL390334.1*) and an additional 11 without names. (Suppl. Fig 1, 3) Overlap between the databases were compared, and replicate entries were removed. The data nomenclature and genomic location was updated and missing values (for example if pseudogene 2 and 3 were identified but 1 was missing) were completed from NCBI (101) and Ensembl (102) databases. Data filtration was performed in multiple steps, based on translation of Ub throughout the open reading frame, and on the presence of evidence at the protein level. For potential protein sequence, the ExPASy (103) *in silico* translation tool was used to accept pseudogene that could translate into a full length Ub protein, as well as the presence of a C-terminal Gly-Gly motif (participating in substrate binding). Expression of Ub pseudogenes with a Gly-Gly motif and at least 90% similarity with the parental Ub protein were then further confirmed by searching for ribosome binding evidence using GWIPS-viz (57,104) database. As an additional step, OpenProt (60,105) and PepQuery (61,62) databases was also searched with the selected pseudogenes for evidence of protein detection in mass spectrometry experiments.

### Sequencing of RPS27AP5 from genomic DNA and mRNA

Genomic DNA was extracted from HeLa cells. Cell pellets were resuspended in 500 µl TAIL buffer (100mM Tris pH8.0, 5mM EDTA, 0.2% SDS, 200mM NaCl) supplemented with 0.1 mg/ml Proteinase K (Roche, Switzerland) and incubated at 37°C overnight. The next day samples were incubated at room temperature with shaking at 1400rpm for 5 min. NaCl was added to the samples to a final concentration of 1.6M and incubated for a further 5 min. Genomic DNA was separated by centrifugation at 13 000rpm at 4°C for 30 min. The supernatant containing the genomic DNA was precipitated by the addition of 700µl of isopropanol. The DNA was pelleted at 13 300 rpm at 4°C for 10 min. The pellet was washed with 70% ethanol, followed by centrifugation at 13 300 rpm at room temperature for 5 min. After the ethanol was removed, the pellet was air dried and then resuspended in DNA Elution Buffer (Monarch DNA Gel Extraction Kit, NEB, USA). Since the *RPS27AP5* sequence, unlike RPS27A, does not contain introns, primers specific to *RPS27A* and *RPS27AP5* common regions were used to amplify *RPS27AP5* gDNA (RPS27A/P5_Seq_FW: 5’-GTTTATTCAGCTTTTCGATCCG-3’ and RPS27A/P5_Seq_RV: 5’-GGCAACTAATTTTGCCATTCTC-3’). The PCR products were run on a 1% agarose gel, and areas corresponding to the expected size were excised and purified from the gel using a gel extraction kit (Monarch DNA Gel Extraction Kit, NEB). Following purification, the PCR products were sent directly for sequencing (Université Laval Sequencing Platform).

RNA was isolated from HeLa cells using TRIzol reagent (Invitrogen, USA) and was reverse transcribed using ProtoScript II reverse transcriptase (New England Biolabs, USA) with primers specific to RPS27A and RPS27AP5 common regions previously used for gDNA amplification. The reaction was incubated at 42°C for 1 h. cDNA specific to *RPS27AP5* was amplified using primers specific to RPS27AP5 (RPS27AP5_FW: 5’-GTTGAACTCTCGGATACAATAGAT-3’ and RPS27AP5_RV: 5’-CATCCACCTTATAATATTTCAGGAG-3’) with an expected product size of 280 base pairs. After PCR amplification using Taq polymerase, samples were run on a 1% agarose gel and areas corresponding to the expected size were excised and purified from the gel using a gel extraction kit (Monarch DNA Gel Extraction Kit, NEB, USA). This PCR product was used for a second round of amplification followed by gel purification. Purified PCR products were inserted into pGEM-T Easy vector (Promega,USA).

### Cloning and plasmids

HA-UBBP3, HA-UBA52P5, HA-RPS27AP1, HA-RPS27A P5 and HA-RPS27AP11 were synthesized as gBlocks containing a stop codon at the end (Integrated DNA technologies, USA) and were inserted first into into pDONR221 plasmid (Life Technologies, Canada) and then using the Gateway cloning system (Life Technologies, Canada) into the pDEST47-3Myc plasmid. For C-terminal epitope labelling of RPS27AP5, the stop codon of the gene was removed by PCR mutagenesis using RPS27AP5_no stop_FW: 5’-CACCCAGCTTTCTTGTACAAAGTTG-3’ and RPS27AP5_no stop_RV: 5’-CTTGTCTTCTGGTTTGTTGAAACAGTAAG-3’ primers. The HA-RPS27A, chimeric sequence containing either wild type Ub and RPS27AP5 coding ribosomal sequences (HA-RPS27A-Ub-RPS27AP5-ribo-Myc) or the reverse (HA-RPS27AP5-Ub-RPS27A-ribo-Myc) were synthesized as gBlocks without a stop codon (Integrated DNA technologies, USA) and were inserted first into a pDONR221 plasmid (Life Technologies, Canada) and then using the Gateway cloning system (Life Technologies, Canada, Canada), into pDEST47-3Myc plasmid (generated by the replacement of GFP in pDEST47).

### Cell culture, transfection and immunoblotting

HeLa cells (ATCC, USA) were grown in Dulbecco’s Modified Eagle Medium (DMEM), and hTERT-RPE-1 cells (ATCC, USA) were grown in Dulbecco’s Modified Eagle Medium/ Nutrient Mixture F-12 (DMEM/F-12) (Life Technologies, Canada), supplemented with 10% fetal bovine serum (FBS) (Invitrogen), 100U/ml penicillin/streptomycin (Wisent Bioprodudcts, Canada), and 2.5µg/ml Amphotericine B (Wisent Bioprodudcts, Canada).

Cells were transfected with control plasmid pcDNA3.1, or plasmid constructs described previously using Lipofectamine 2000 or Lipofectamine LTX reagent according to the manufacturer’s instructions (Thermo Fisher Scientific, USA). For MG132 treatment for western blot analysis, cells were incubated in culture media containing a 20 µM final concentration of MG132 (Sigma-Aldrich, USA) for 5 h. 48 hours after transfection, cells were lysed in 1x Laemmli solution, and proteins were separated on SDS-PAGE gel. Proteins were transferred to nitrocellulose membrane and immunoblotting was performed with the following antibodies: HA (#26183, 2-2.2.14, Invitrogen, USA, 1:1000), Myc (#2278, 71D10, Cell Signaling, USA, 1:400), RPS3 (#376008, Santa Cruz Biotechnology, USA, 1:1000), RPL11 (#79352, Abcam, USA, 1:1000), H3K27me3 (#9733, Cell Signaling, 1:1000), p15/p16 (#377412, Santa Cruz Biotechnology, 1:200) and GAPDH (#5174S, Cell Signaling, USA, 1:50000).

### Immunofluorescence

Cells were seeded onto glass coverslips in a 24-well cell culture plate to reach 50% confluency the next day. Cells were transfected with plasmids containing HA epitope tagged Ub, Ub^KEKS^, UBBP3, UBA52P6, RPS27AP1, RPA27AP5 and RPS27AP11 or with HA-RPS27A-Myc, HA-RPS27AP5-Myc epitope tagged plasmids using JetPrime (Polyplus-transfection SA, France) reagent. For MG132 treatment, cells were incubated in culture media containing a 1.2 µM final concentration of MG132 for 24 hours. 48 hours after transfection, cells were washed twice with 1x PBS, fixed and permeabilised with 100% ice cold methanol. Coverslips were blocked on ice with 1% BSA in 1x PBS for 30 minutes followed by incubation overnight at 4°C with anti HA-488 antibody (#26183, 16B12, Invitrogen, USA, 1:1000) and where applicable with Myc (#2278, 71D10, Cell Signaling, USA, 1:400) antibody. The following day, cells were washed twice with 1x PBS followed by incubation with DAPI solution (1µg/µl) for 8 minutes in 1x PBS and washed again twice. Coverslips also probed with Myc antibody were washed three times with ice cold 1xPBS and incubated with AlexaFluor 546 (#11010, Invitrogen, USA, 1:800) secondary antibodies in 10% goat serum in 1xPBS at 4°C for 1 h prior to incubation with a DAPI solution. Coverslips were mounted on microscope slides with Immuno-mount (Thermo Fisher Scientific, USA) mounting media and stored at 4°C in the dark until imaging. Images were acquired on a Zeiss LSM 880 confocal microscope using a 40× 1.4NA plan Apo objective.

### Co-immunoprecipitation

HeLa cells were transfected with HA-RPS27A-Myc or HA-RPS27AP5-Myc constructs using Lipofectamine LTX reagent (Thermo Fisher Scientific, USA), and were incubated for 48 h. Cells were washed with 1xPBS, harvested by scraping and centrifuged at 1500rpm for 5 min at 4°C. Pelleted cells were resuspended in Lysis buffer (50mM Tris pH 7.4, 150mM NaCl, 5mM MgCl_2_ 1% Triton X-100, protease inhibitors (Roche Complete Inhibitor cocktail, Roche, Switzerland)) and incubated for 20 min at 4°C on a rotator. Lysates were cleared by centrifugation at 10 000rpm for 10 min at 4°C. Equal concentration of total protein extracts were incubated with Myc (#1-9E10.2 hybridoma supernatant, ATCC, USA) or HA (#11583816001, 12CA5, Sigma-Aldrich, USA) antibodies for 2h at 4°C on a rotator. Antibody binding was followed by incubation with Protein A/G magnetic beads (Thermo Fisher Scientific, USA) for 1h at 4°C on a rotator. Beads were then washed with Lysis buffer three times, and samples were processed for on-beads digestion.

### On-beads digestion

Samples were washed five times in 20mM ammonium bicarbonate solution (NH_4_HCO_3_ in LC/MS grade H_2_O). After the washes, samples were reduced at 60°C for 30 minutes using 10mM DTT (dithioreitol) followed by alkylation at room temperature for 1 hour using 15 mM of 2-chloroacetamide. Proteins were digested with 1µg trypsin (Trypsin Gold, Promega Corporation) overnight at 37°C in a shaker at 1250 rpm. Tryptic peptides were extracted by the addition of 1% formic acid (FA) (Fisher Chemical, USA), followed by further two incubations of the beads with 60% acetonitrile (CH_3_CN)/ 1% FA solution in LC/MS grade H_2_O. The extraction solutions containing the digested peptides were pooled and the peptides were dried in a speedvac and resuspended in 0.1% TFA (trifluoroacetic acid) in LC/MS grade H_2_O. Desalting was performed using C-18 tips (Thermo Fisher Scientific, USA), equilibrated with three washes in 100% CH_3_CN and 0.1% TFA in LC/MS grade H_2_O. Peptides were bound to the column by pipetting the samples up and down. After three washes of 0.1% TFA, elution was performed using Elution buffer (1% FA, 50% CH_3_CN in LC/MS grade H_2_O). The eluted peptides were dried in a speedvac and resuspended in 1% FA in LC/MS grade H_2_O to be processed by mass spectrometry analysis.

### Mass spectrometry analysis

Digested peptides were injected into an HPLC (nanoElute, Bruker Daltonics) and loaded onto a trap column with a constant flow of 4 µl/min (Acclaim PepMap100 C18 column, 0.3 mm id x 5 mm, Dionex Corporation), then eluted onto an analytical C18 Column (1.9 µm beads size, 75 µm x 25 cm, PepSep). Peptides were eluted over a 2-hour gradient of CH_3_CN (5-37%) in 0.1% FA at 400 nL/min while being injected into a TimsTOF Pro ion mobility mass spectrometer equipped with a Captive Spray nano electrospray source (Bruker Daltonics). Data was acquired using diaPASEF mode, where for each single TIMS (100 ms), 1 mobility window consisting of 27 mass steps (m/z between 114 to 1414 with a mass width of 50 Da) per cycle (1.27 seconds duty cycle) was used. These steps covered the diagonal scan line for +2 and +3 charged peptides in the m/z-ion mobility plane. Identification and quantification of proteins was performed using the MaxQuant software (version 2.0.3.0) (106,107). Statistical analysis was performed using Prostar software (v1.26.3, DAPAR v1.26.1) (108,109). Data filtering was performed by removing missing POV (partially observed values) data <= 1 in each condition, proteins with <= 1 unique peptide, as well as data labelled as ‘Reverse’, ‘Potential_contaminant’, and ‘Only_identified_by_site’. For normalization, Quantile Centering within conditions was used (Quantile: 0.15). POV imputation: slsa, MEC (missing on entire condition) imputation: detQuantile (Quantile: 0.5, Factor: 0.2). Limma test was used for hypothesis testing with logFC threshold: 1 and −log10 p value: 1.3.

### Luciferase assay

Hela cells were grown in 12-well plates and co-transfected with 400ng of pIRES-p53, pIRES-VEGFA, pIRES-VEGFB, pIRES-FGF1 or pIRES-FGF2 (110–112) and 400ng of pDEST47-3Myc-RPS27A or pDEST47-3Myc-RPS27AP5 using JetPrime (Polyplus-transfection SA, France). After 48h, Luciferase assay was performed on a LUMIstar luminometer (BMG Labtech, USA) by using Dual Luciferase-Reporter Assay System (Promega, USA) according to the manufacturer’s instructions. Firefly luciferase activity were normalized to Renilla luciferase activity.

### Ribosome extraction

HeLa cells were transfected with HA-RPS27A-Myc or HARPS27AP5-Myc constructs using Lipofectamine LTX reagent (Thermo Fisher Scientific, USA), and were incubated for 48 h. Cells were then treated with 100 µg/ml of cycloheximide (CHX) (Thermo Fisher Scientific, USA) for 5 min at 37°C. Cells were then washed with ice cold 1xPBS supplemented with 100 µg/ml CHX. After harvesting them by scraping in the same solution used for washing, cells were pelleted at 1500rpm for 5 min at 4°C. Cell pellets were then resuspended in Ribosome Lysis Buffer (20mM Tris pH7.4, 10mM MgCl_2_, 300mM KCl, 1mM DTT, 40U/ml RiboLock RNase inhibitor (Thermo Fisher Scientific, USA), 1% Triton X-100, 100 µg/ml CHX) and incubated on ice for 20 min. Cell lysates were cleared by centrifugation at 10 000 rpm for 10 min at 4°C. The cleared lysate was then layered over a Sucrose solution (20mM Tris pH7.4, 10mM MgCl_2_, 300mM KCl, 20% sucrose, 100 µg/ml CHX). Ribosomes were separated by centrifugation at 70 000 rpm at 4°C in a TLA-110 ultracentrifuge rotor for 4 h 30 min. Ribosome pellets were resuspended in SDS-PAGE loading buffer and proteins were separated on an SDS-PAGE gel.

### Polysome profiling

HeLa cells were transfected with HA-RPS27A-Myc or HA-RPS27AP5-Myc constructs and were incubated for 48 h. Cells were then treated with 100 µg/ml of cycloheximide (CHX) (Thermo Fisher Scientific, USA) for 5 min at 37°C. Cells were then washed three times with ice cold 1xPBS supplemented with 100 µg/ml CHX and were harvested by scraping in the same solution. Harvested cells were pelleted at 2500rpm for 10 min at 4°C. Samples were lysed using Ribosome Lysis Buffer and after incubation at 4°C for 20 min, protein extracts were cleared by centrifugation at 13 000rpm for 10 min at 4°C. 3-5mg of total protein extracts were loaded onto a 10-45% sucrose gradient in polyallomer tubes (14×89 mm, Setton). Samples were centrifuged for 3 h at 40 000 rpm in a SW41swing out rotor (Beckman Coulter). The separated gradient was then fractionated by upward displacement with 55% sucrose solution using a gradient fractionator (Brandel Inc., USA) connected to a UA-6 UV monitor (Teledyne Isco, USA) for continuous measurement of the absorbance at 254 nm. Fractions of 650µl were collected and proteins were precipitated using trichloroacetic acid (TCA) (Thermo Fisher Scientific, USA) precipitation, and pellets were resuspended in SDS-PAGE loading buffer. Ribosome distribution profiles were analysed using a DI-1100 instrument and the WinDaq software (DATAQ Instruments, v1.61).

### Translating ribosome bound RNA pulldown (TRAP) assay

HeLa cells were transfected with HA-RPS27A-Myc or HA-RPS27AP5-Myc constructs and were incubated for 48 h. Cells were treated with cycloheximide (CHX) (Thermo Fisher Scientific, USA), harvested and lysed in Ribosome Lysis Buffer as described above. 1.5 mg of total protein lysate was incubated first with empty magnetic beads for 30 min, followed by incubation with MycTrap beads (Chromotek, Germany) for 1h at 4°C. Beads were washed three times prior to the addition of cell lysates with 0.15 M KCl Wash Buffer (10mM HEPES, pH7.2-7.5, 5mM MgCl_2_, 150mM KCl, 1% NP-40). In addition, MycTrap beads were pre-incubated with yeast tRNA (Roche, Switzerland) to block non-specific RNA binding in 0.15M KCl Wash Buffer supplemented with 0.5mM DTT and 40U/ml RiboLock RNase inhibitor (Thermo Fisher Scientific, USA). Final washes were performed three times in 0.35M KCl Wash Buffer (10mM HEPES, pH7.2-7.5, 5mM MgCl_2_, 350mM KCl, 1% NP-40). Beads were then resuspended in 300 µl RLT buffer (Qiagen RNeasy kit, Qiagen, Germany) and incubated at room temperature for 5 min. The supernatant was saved and recovered RNA was cleaned using Qiagen RNeasy kit (Qiagen, Germany).

### Preparation and sequencing of total RNAseq libraries

Each of the RNAseq libraries was generated from 50 ng of total RNA free of genomic DNA using the NEBNext Single Cell/Low input RNA Library Prep kit for Illumina (E6420S) and following the protocol for Low input. The resulting libraries were subjected to a total of 11 amplification cycles and then purified using Ampure XP 0.9X beads. Quality and size were assessed with an Agilent 2100 Bioanalyzer. Libraries were then quantified using a Qubit fluorometer, pooled at an equimolar concentration and diluted to 1.8 pM then sequenced on Illumina’s NextSeq 500 using a NextSeq 500/550 High Output kit v2.5 (75 cycles) 2×40bp.

### RNA-seq analysis

Reads were trimmed using fastp v0.23.2 (113) with the following parameters “-- qualified_quality_phred 30 --length_required 20 --cut_window_size 1 --cut_mean_quality 30 --cut_front --cut_tail” to remove adapters and low-quality reads. The quality of the reads was assessed before and after trimming using FastQC v0.11.5. The human transcriptome was generated from the human genome (GRCh38 assembly, v107 in Ensembl) using gffread (available from Cufflinks v2.2.1 (114)). An index was created from this transcriptome using the “index” mode of Kallisto v0.48.3 (115) and the following parameter “--kmer-size=31”. Trimmed reads were then pseudoaligned to this human transcriptome and quantified using the “quant” mode of Kallisto with the following parameters “--bias --bootstrap-samples=50”. Differential expression analyses were finally conducted at the gene-level using tximport v1.22.0 (116) and DESeq2 v1.34.0 (117). All of the analyses to generate the abundance datasets were organized in a Snakemake workflow available at Github (https://github.com/etiennefc/RPG_pseudogenes_RNA_Seq.git). For the volcano plot, only protein coding genes were visualised using the non-adjusted p-value. Results were visualised on Graphpad Prism 9 v9.5.1 (733).

### CUT and RUN and qPCR analysis

Cleaveage Under Targets and Release Using Nuclease experiment was performed using a CUT & RUN Assay Kit (Cell Signaling, USA), according to the manufacturer’s instructions. Briefly, HeLa cells were seeded in a 60 mm dish and were transfected with HA-RPS27A-Myc or HA-RPS27AP5-Myc constructs, and were incubated for 48 h. For negative controls, untransfected cells were used. On the day of the assay, cells were trypsinised, and for each sample 100 000 cells were counted using a hemocytometer (for each reaction as well as for input). After washing, the harvested cells were bound to Concavalin A-coated magnetic beads, and subsequently permeabilised using digitonin. Primary antibody binding was then performed at 4°C for 2 h. For negative control rabbit IgG (DA1E) (#66362, Cell Signaling, USA), for positive control H3K4me3 (C42D8) (#9751, Cell Signaling, USA) and for the experimental reaction H3K27me3 (C36B11) (#9733, Cell Signaling, USA) antibodies were used. After washing the cells, a pAG-MNase (proteinA/G-Micrococcal Nuclease) fusion binding to the primary antibody heavy chain was performed at 4°C for 1h. Cells were then washed again, and pAG-MNase was activated by the addition of cold CaCl_2_. Site-specific cleavage was performed at 4°C for 30 min, and the digestion was stopped by the addition of 1x Stop Buffer containing spike-in DNA for normalization. Cleaved DNA fragments were released by incubating samples at 37°C for 10 min followed by centrifugation at 4°C for 2 min at 16 000g. The supernatant containing the enriched chromatin samples was saved and purified using DNA Purification Buffers and Spin Columns (#14209, Cell Signaling, USA). The recovered chromatin samples were then subjected to qPCR analysis using primer pairs specific to *INK4A* promoter region sequences (INK4A−1kb_FW: CTCAAAGCGGATAATTCAAGAGC, INK4A−1kb_RV: AAGCCTTAAGAACAGTGCCACAC, INK4A−0.2kb_FW: ACCCCGATTCAATTTGGCAG, INK4A−0.2kb_RV: AAAAAGAAATCCGCCCCCG) (118).

For the qPCR assays, primers were individually resuspended to 20–100 μM stock solutions in Tris-EDTA buffer and diluted as a primer pair to 1 μM in RNase DNase-free water. qPCRs were performed in 10 μL in 96-well plates on a CFX OPUS-96 thermocycler (Bio-Rad) with 5 μL of 2X PerfeCTa SYBR Green Supermix (Quantabio), 3 μL of DNA (1:2 dilution) and 200 nM (final concentration; 2 μL) primer pair solutions. The following cycling conditions were used: 3 min at 95°C and 50 cycles of 15 s at 95°C, 30 s at 60°C, and 30 s at 72°C. Relative expression levels were calculated using the qBASE framework (119) using the primer pair provided by the manufacturer for the spike-in DNA (*E. coli* DNA (5ng)) as housekeeping genes for normalization. Primer design and validation were evaluated as described elsewhere (120). In every qPCR run, a no-template control was performed for each primer pair, and these were consistently negative. All qPCR assays were performed by the RNomics Platform of the Université de Sherbrooke.

## Acknowledgements

We thank Frédéric Catez and Anne-Catherine Prats (CRCL, Lyon) for the IRES luciferase plasmids, Dr. Anne-Marie Landry-Voyer for technical advice with ribosome extraction and polysome profiling. Funding was provided from the Canadian Institutes for Health Research, grant number #398925 to FMB. FMB is a FRQS Senior scholar (award number #281824). XR is a recipient of a Canada Research Chair in Functional Proteomics and Discovery of Novel Proteins. XR, FMB and MSS are members of the FRQS-funded “Centre de Recherche du CHUS”.

## Author contributions

AM.: Conceptualization, analysis, investigation, visualization, writing original draft. DL: Mass spectrometry run and analysis. JR: Luciferase assay setup. EFC: Analysis of TRAP RNA-seq data. FMB and XR.: Conceptualization, supervision, funding acquisition, editing original draft.

## Data availability

The mass spectrometry proteomics data have been deposited to the ProteomeXchange Consortium via the PRIDE partner repository with the dataset identifier PXD047544. The RNAseq data has been deposited at NCBI GEO database, accession GSE24489245.

---

**Supplementary Figure 1.**
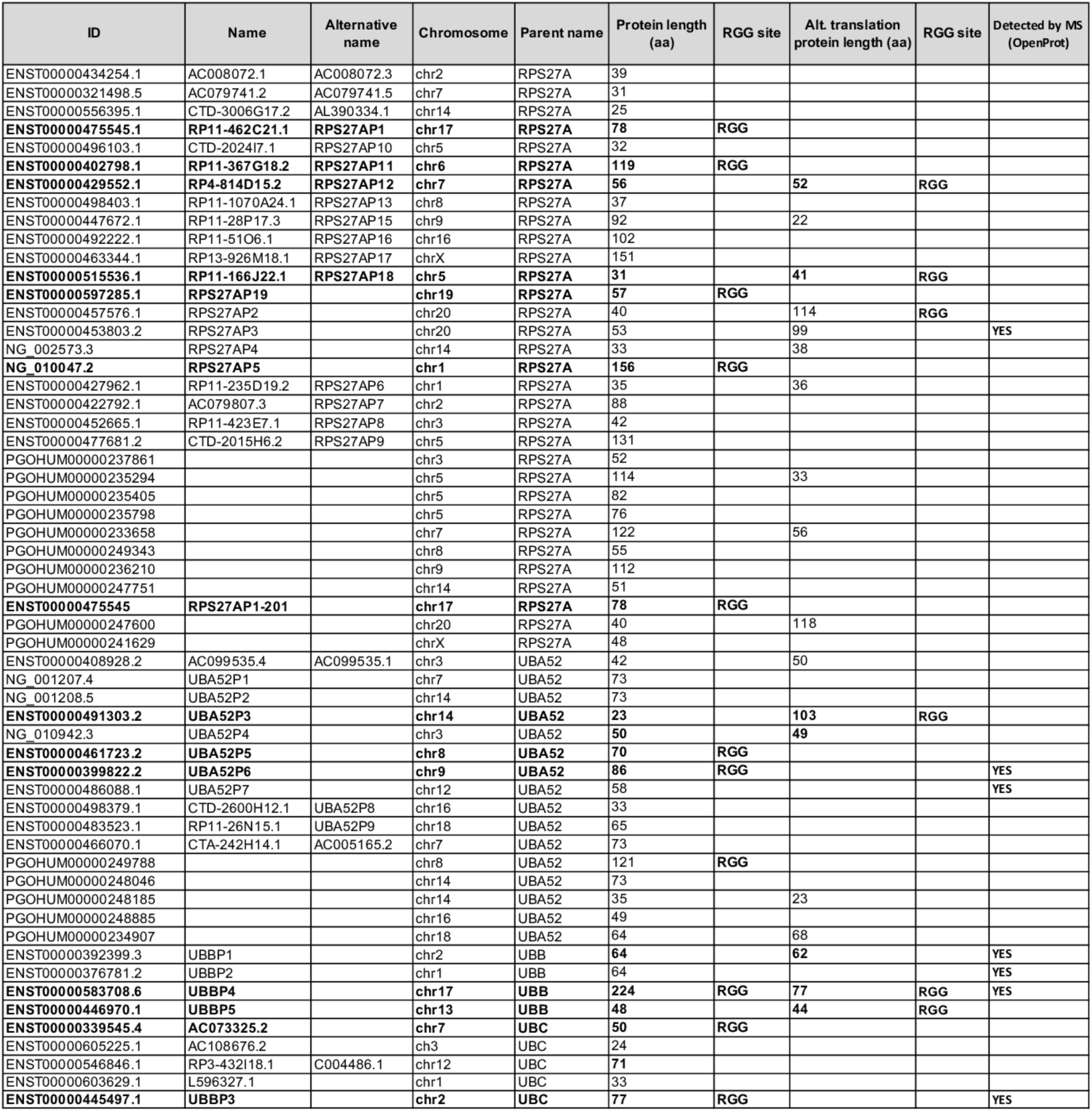
List of collected Ub pseudogenes. List of collected Ubl pseudogenes from Pseudogene.org (GENCODE v.10) and Pseudofam/PseudoPipe with locations, parental genes and Gly-Gly motifs after *in silico* translation.

**Supplementary Figure 2.**
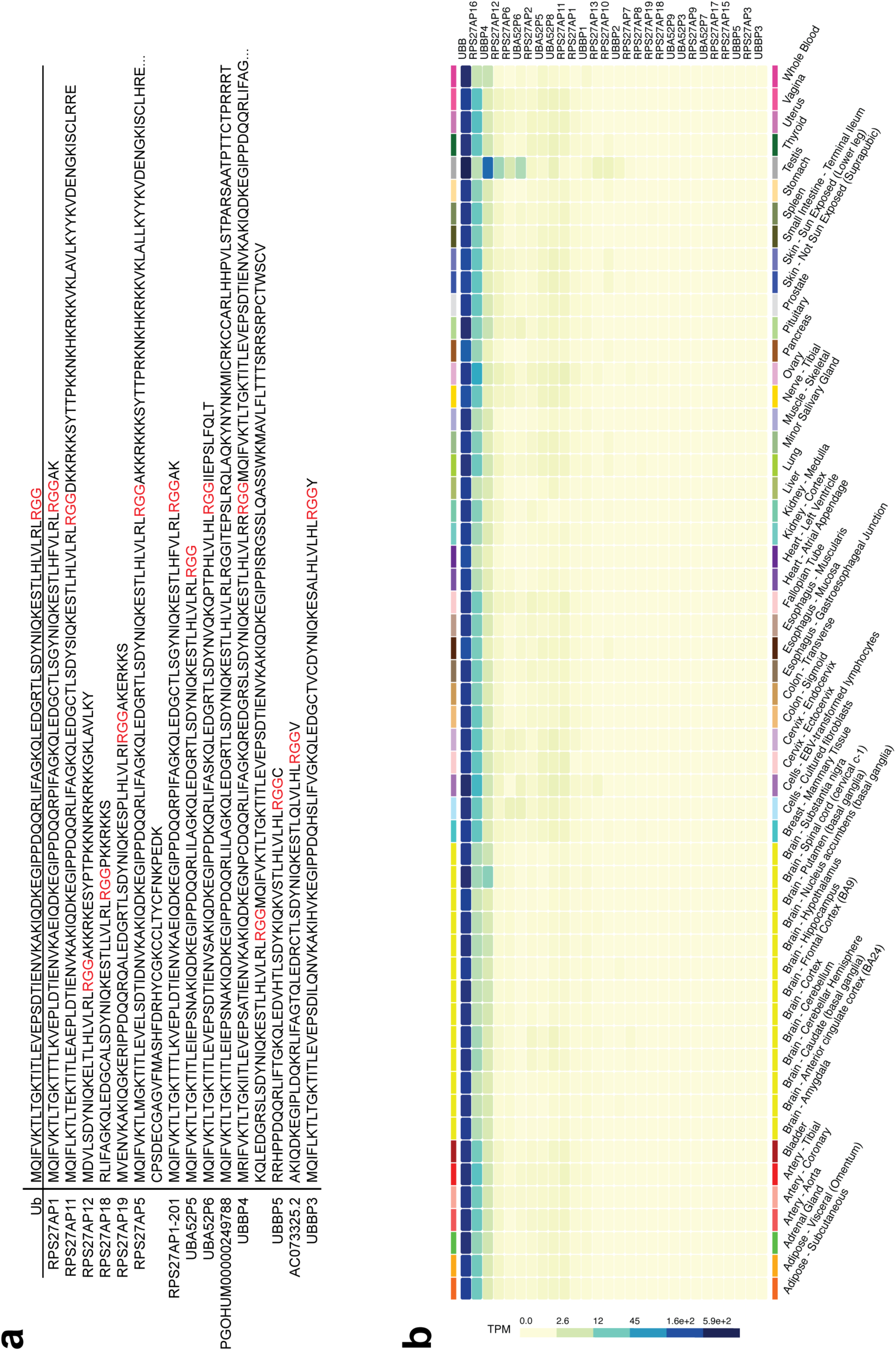
Ub pseudogenes with gene expression levels and translated protein sequences. A. Translated sequences of Ub pseudogenes with RGG motifs. B. Expression levels analysis of the collected Ub pseudogenes from the Genotype-Tissue Expression (GTEx) Portal (121) (only ones with gene names were included in the search). Transcript expression is displayed as transcripts per million (TPM).

**Supplementary Figure 3.**
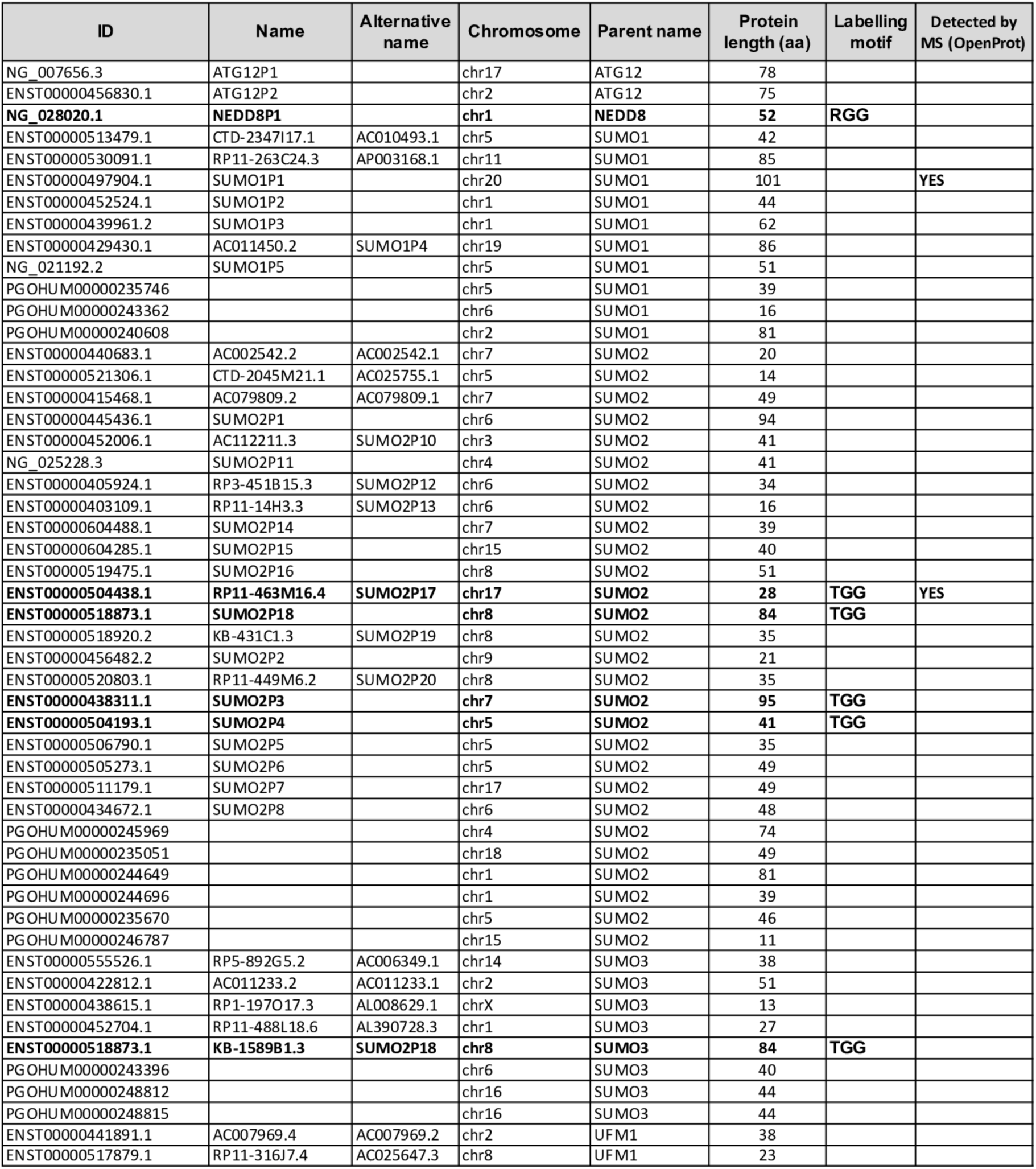
List of collected Ubl pseudogenes. List of collected Ubl pseudogenes from Pseudogene.org (GENCODE v.10) and Pseudofam/PseudoPipe with locations, parental genes and Gly-Gly motifs after *in silico* translation.

**Supplementary Figure 4.**
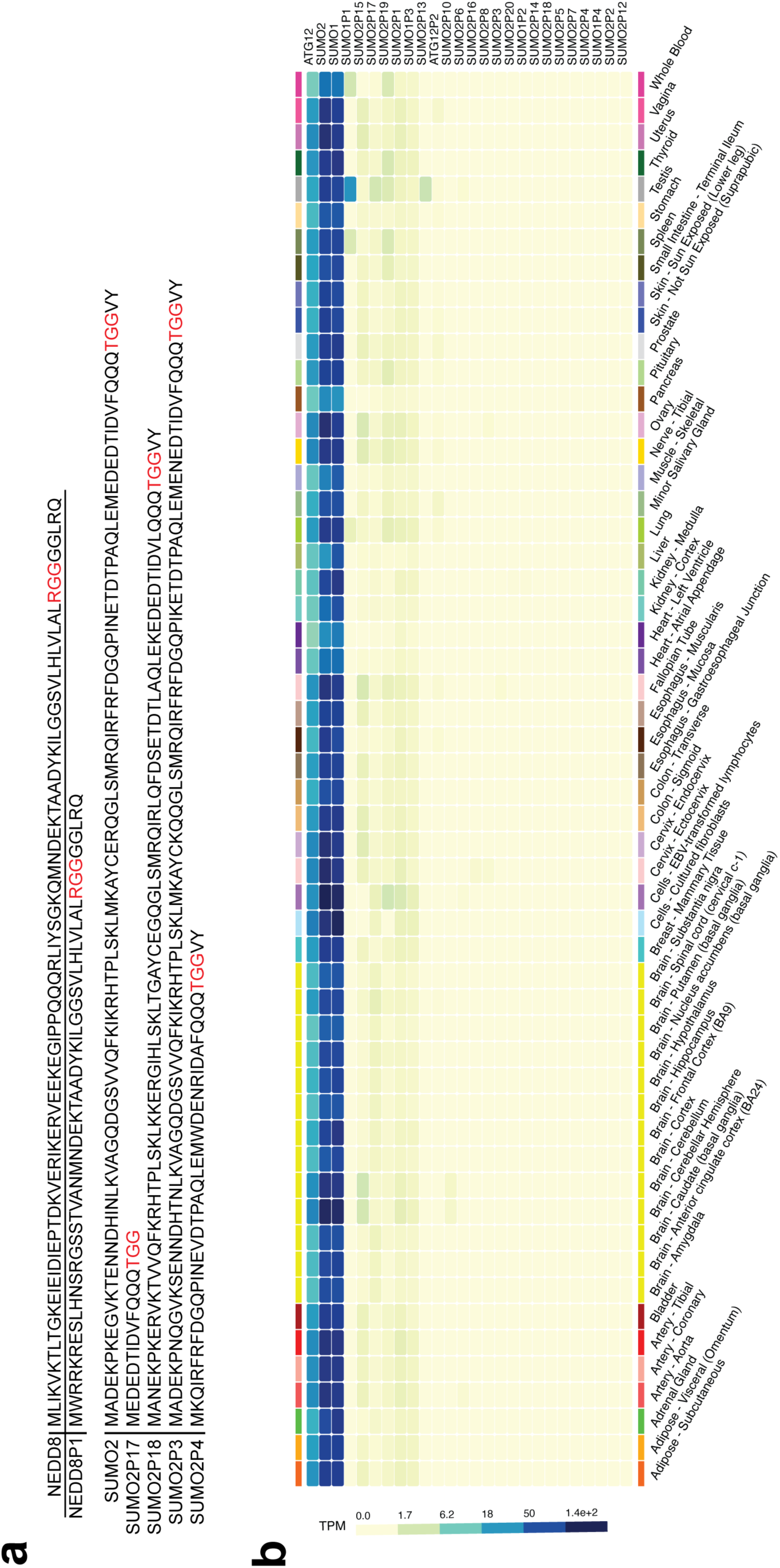
Ubl pseudogenes with gene expression levels and translated protein sequences. A. Translated sequence of Ubl pseudogenes with RGG or TGG motifs. B. Expression levels analysis of the collected Ubl pseudogenes from the Genotype-Tissue Expression (GTEx) Portal (121) (only ones with gene names were included in the search). Transcript expression is displayed as transcripts per million (TPM).

**Supplementary Figure 5.**
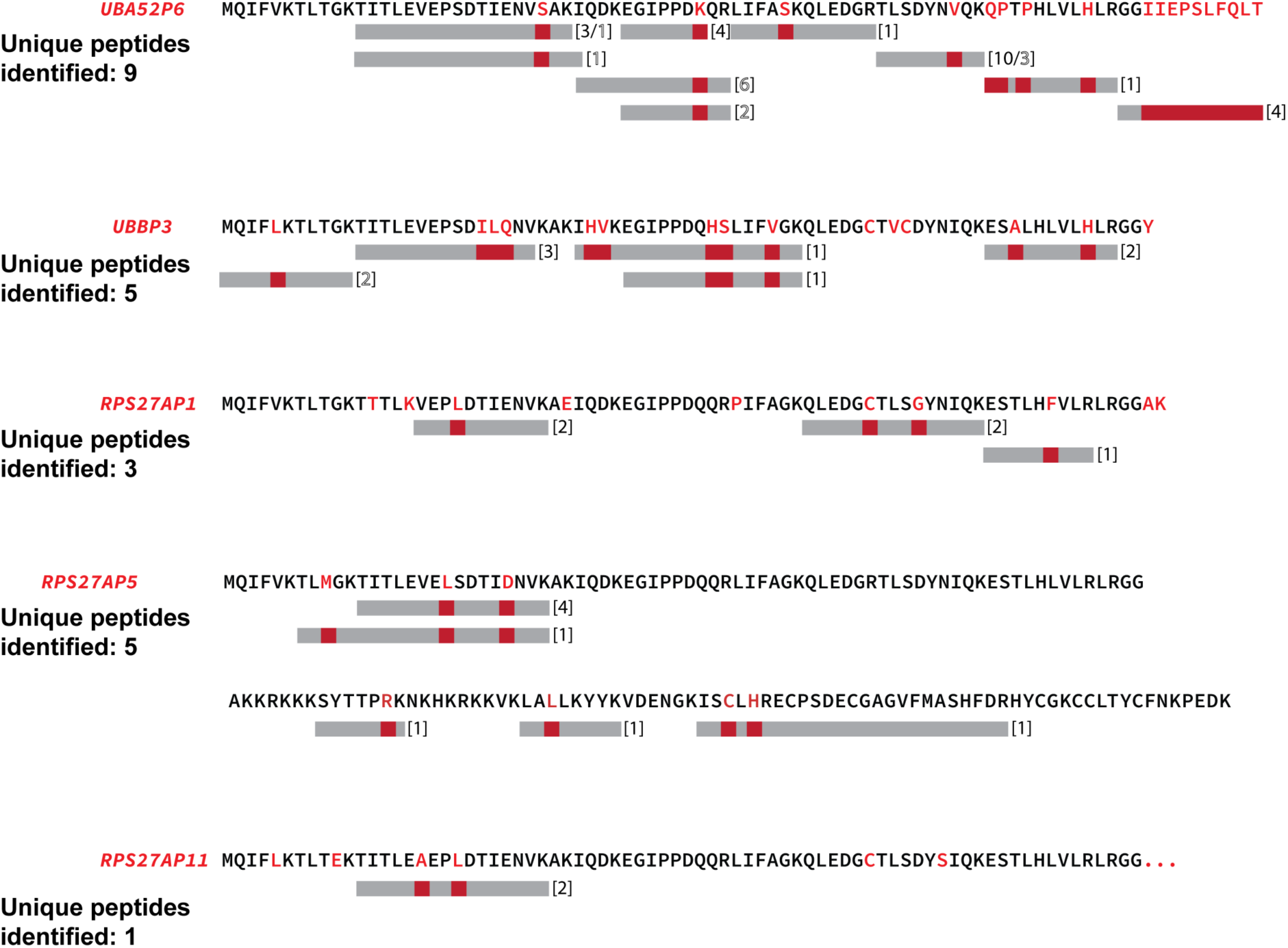
Protein expression evidence of selected candidate pseudogenes. Identified unique peptides for each gene from PepQuery and OpenProt databases. The numbers in bracket represents the number of individual studies in which the peptides were found. Black numbers correspond to results obtained from PepQuery while black outlined numbers correspond to data obtained from OpenProt.

**Supplementary Figure 6.**
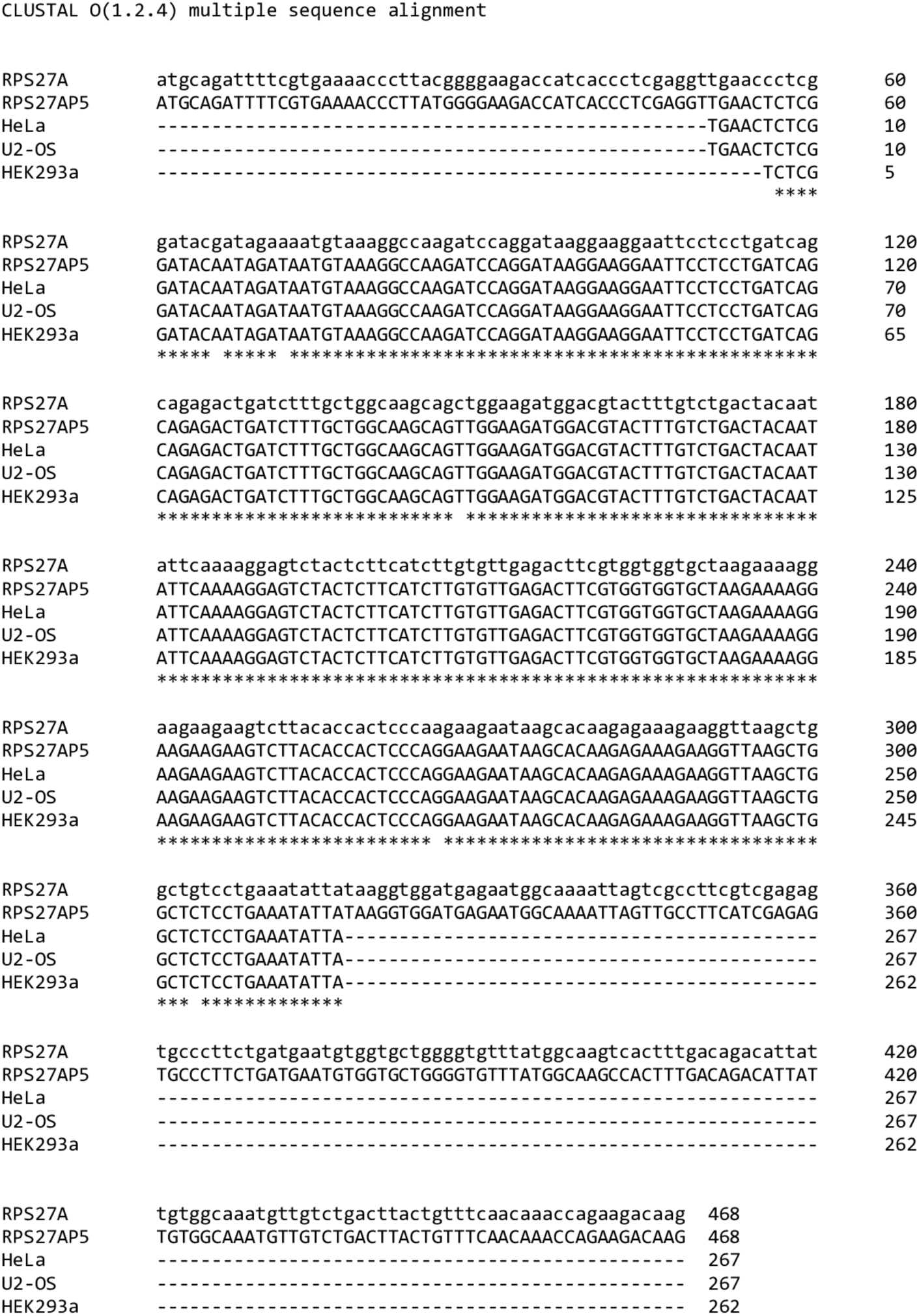
Clustal alignment of sequencing results. RPS27AP5 specific PCR amplification and sequencing results in three different cell lines (HeLa, U2-OS, HEK293a) aligned with the reference sequence of RPS27AP5 and RPS27A using the Clustal tool.

**Supplementary Figure 7.**
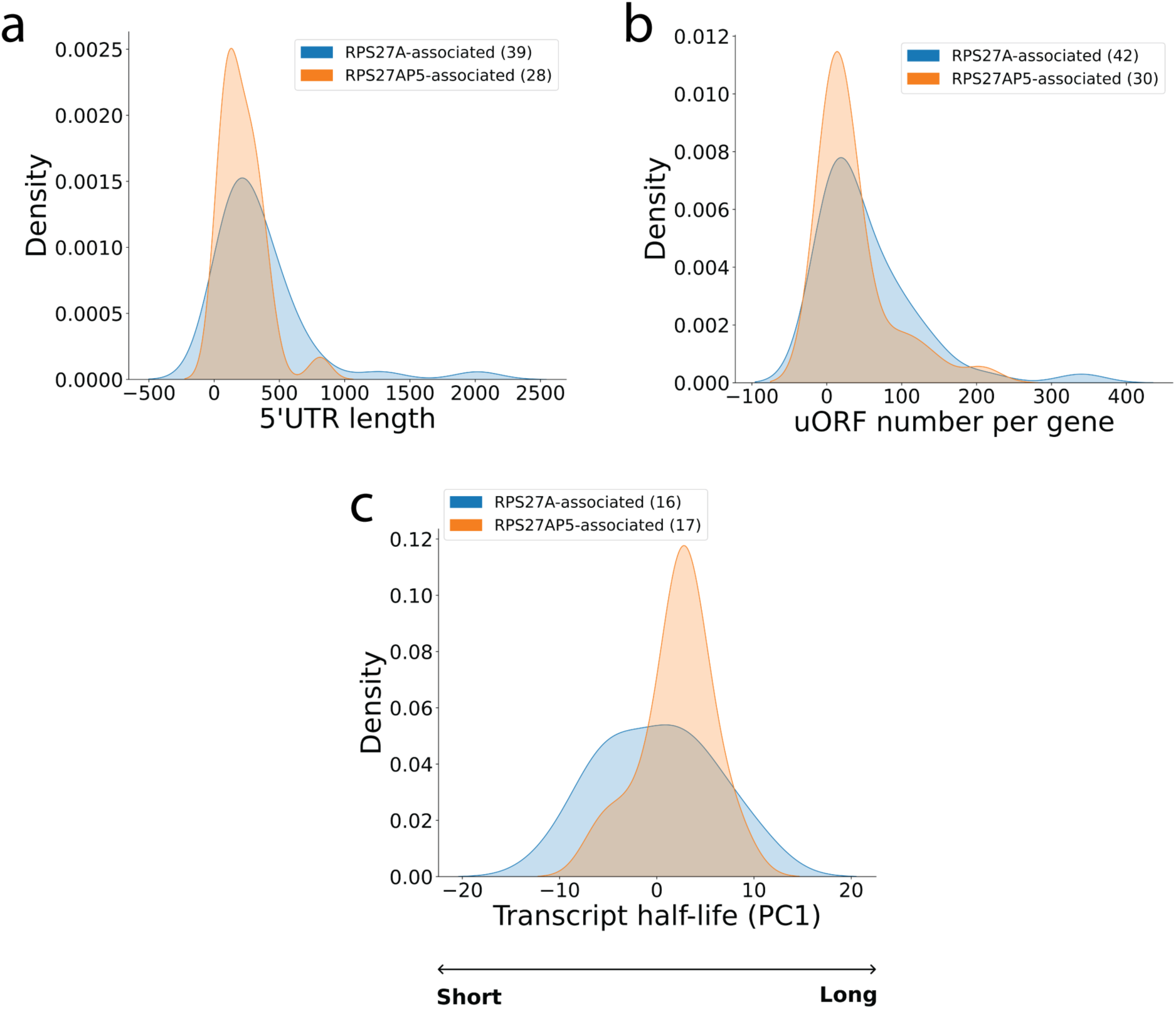
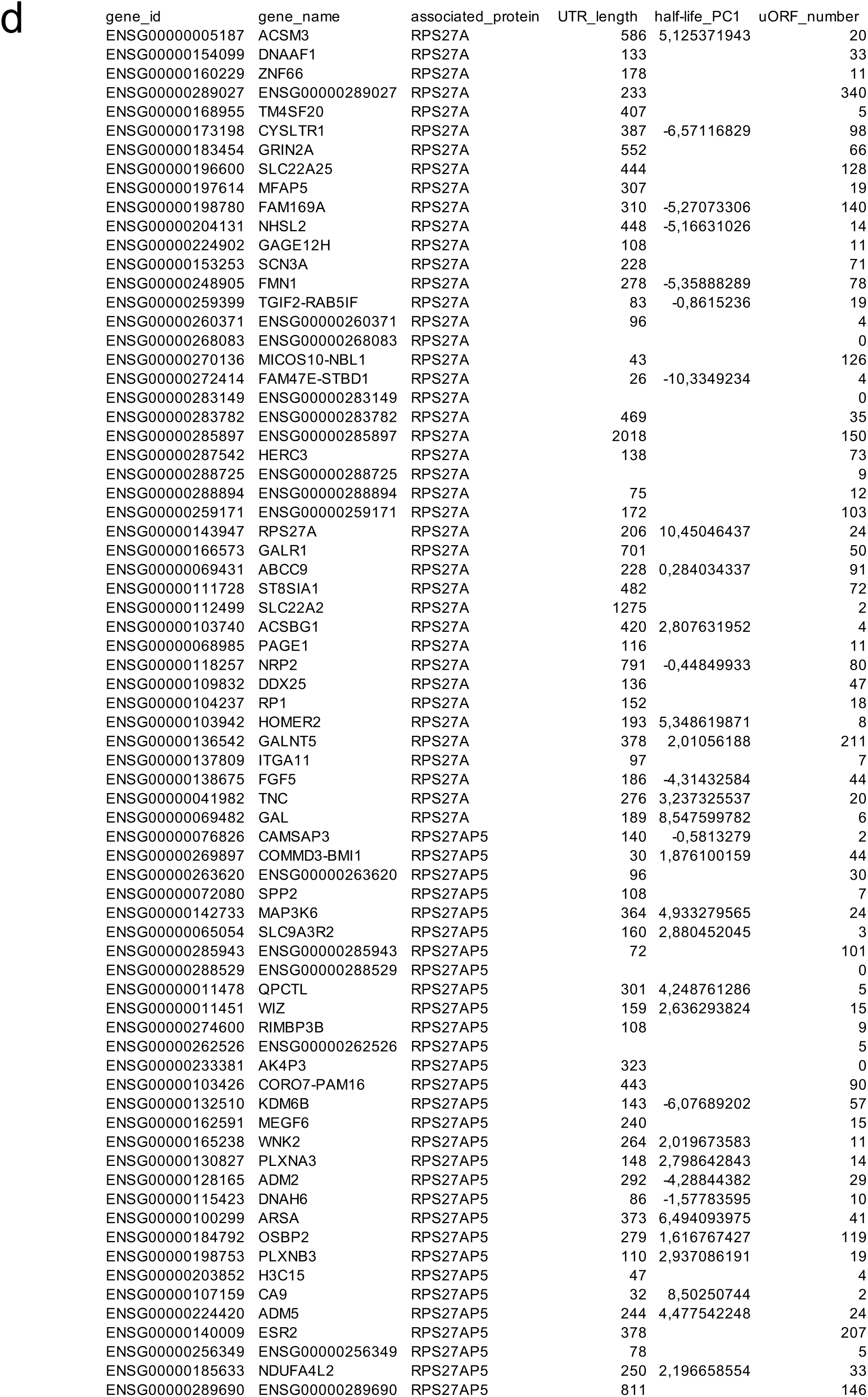
Feature analyses of mRNAs found interacting with ribosomes containing either S27a or S27aP5. A. 5’ UTR length (bp) of mRNAs found interacting with ribosomes containing either S27a or S27aP5. B. Number of upstream open reading frames (uORFs) per of mRNAs found interacting with ribosomes containing either S27a or S27aP5. C. Half-life of mRNAs found interacting with ribosomes containing either S27a or S27aP5. Values are represented as principal component analysis (PCA)-derived PC1 value.

## Supplementary tables

**Supplementary Table 1 – Extended data table of collected and curated Ubiquitin pseudogenes.**

**Supplementary table 2 – Extended data table of collected and curated Ubiquitin-like pseudogenes.**

**Supplementary table 3 – Proteomics data for the comparison of Ub and UbP5 interactomes.**

**Supplementary table 4 – Proteomics data for the comparison of S27a and S27aP5 interactomes.**

**Supplementary table 5 –RNA-Seq data of protein-coding RNAs identified from the TRAP assay using either Myc-RPS27a or Myc-RPS27aP5.**

## References

1. Harrison, P. M., Zheng, D., Zhang, Z., Carriero, N., and Gerstein, M. (2005) Transcribed processed pseudogenes in the human genome: an intermediate form of expressed retrosequence lacking protein-coding ability. Nucleic Acids Res 33, 2374–2383

2. Tutar, Y. (2012) Pseudogenes. Comparative and functional genomics 2012, 424526

3. Zhang, Z. D., Frankish, A., Hunt, T., Harrow, J., and Gerstein, M. (2010) Identification and analysis of unitary pseudogenes: historic and contemporary gene losses in humans and other primates. Genome biology 11, R26

4. Mighell, A. J., Smith, N. R., Robinson, P. A., and Markham, A. F. (2000) Vertebrate pseudogenes. Febs Lett 468, 109–114

5. Long, M., Deutsch, M., Wang, W., Betran, E., Brunet, F. G., and Zhang, J. (2003) Origin of new genes: evidence from experimental and computational analyses. Genetica 118, 171–182

6. Esnault, C., Maestre, J., and Heidmann, T. (2000) Human LINE retrotransposons generate processed pseudogenes. Nat Genet 24, 363–367

7. Vanin, E. F. (1984) Processed pseudogenes. Characteristics and evolution. Biochimica et biophysica acta 782, 231–241

8. Ohshima, K., Hattori, M., Yada, T., Gojobori, T., Sakaki, Y., and Okada, N. (2003) Whole-genome screening indicates a possible burst of formation of processed pseudogenes and Alu repeats by particular L1 subfamilies in ancestral primates. Genome biology 4, R74

9. Zhang, Z., Harrison, P. M., Liu, Y., and Gerstein, M. (2003) Millions of years of evolution preserved: a comprehensive catalog of the processed pseudogenes in the human genome. Genome research 13, 2541–2558

10. Zheng, D., Frankish, A., Baertsch, R., Kapranov, P., Reymond, A., Choo, S. W., Lu, Y., Denoeud, F., Antonarakis, S. E., Snyder, M., Ruan, Y., Wei, C. L., Gingeras, T. R., Guigo, R., Harrow, J., and Gerstein, M. B. (2007) Pseudogenes in the ENCODE regions: consensus annotation, analysis of transcription, and evolution. Genome research 17, 839–851

11. Goncalves, I., Duret, L., and Mouchiroud, D. (2000) Nature and structure of human genes that generate retropseudogenes. Genome research 10, 672–678

12. Zhang, Z., Harrison, P., and Gerstein, M. (2002) Identification and analysis of over 2000 ribosomal protein pseudogenes in the human genome. Genome research 12, 1466–1482

13. Maestre, J., Tchenio, T., Dhellin, O., and Heidmann, T. (1995) mRNA retroposition in human cells: processed pseudogene formation. Embo J 14, 6333–6338

14. D’Errico, I., Gadaleta, G., and Saccone, C. (2004) Pseudogenes in metazoa: origin and features. Briefings in functional genomics & proteomics 3, 157–167

15. Xu, J., and Zhang, J. (2016) Are Human Translated Pseudogenes Functional? Molecular biology and evolution 33, 755–760

16. Svensson, O., Arvestad, L., and Lagergren, J. (2006) Genome-wide survey for biologically functional pseudogenes. PLoS computational biology 2, e46

17. Hawkins, P. G., and Morris, K. V. (2010) Transcriptional regulation of Oct4 by a long non-coding RNA antisense to Oct4-pseudogene 5. Transcription 1, 165–175

18. Milligan, M. J., and Lipovich, L. (2014) Pseudogene-derived lncRNAs: emerging regulators of gene expression. Frontiers in genetics 5, 476

19. Watanabe, T., Totoki, Y., Toyoda, A., Kaneda, M., Kuramochi-Miyagawa, S., Obata, Y., Chiba, H., Kohara, Y., Kono, T., Nakano, T., Surani, M. A., Sakaki, Y., and Sasaki, H. (2008) Endogenous siRNAs from naturally formed dsRNAs regulate transcripts in mouse oocytes. Nature 453, 539–543

20. Guo, X., Zhang, Z., Gerstein, M. B., and Zheng, D. (2009) Small RNAs originated from pseudogenes: cis- or trans-acting? PLoS computational biology 5, e1000449

21. Ebert, M. S., and Sharp, P. A. (2010) Emerging roles for natural microRNA sponges. Curr Biol 20, R858–861

22. Lou, W., Ding, B., and Fu, P. (2020) Pseudogene-Derived lncRNAs and Their miRNA Sponging Mechanism in Human Cancer. Frontiers in cell and developmental biology 8, 85

23. Brosch, M., Saunders, G. I., Frankish, A., Collins, M. O., Yu, L., Wright, J., Verstraten, R., Adams, D. J., Harrow, J., Choudhary, J. S., and Hubbard, T. (2011) Shotgun proteomics aids discovery of novel protein-coding genes, alternative splicing, and “resurrected” pseudogenes in the mouse genome. Genome research 21, 756–767

24. Ji, Z., Song, R., Regev, A., and Struhl, K. (2015) Many lncRNAs, 5’UTRs, and pseudogenes are translated and some are likely to express functional proteins. eLife 4, e08890

25. Dubois, M. L., Meller, A., Samandi, S., Brunelle, M., Frion, J., Brunet, M. A., Toupin, A., Beaudoin, M. C., Jacques, J. F., Levesque, D., Scott, M. S., Lavigne, P., Roucou, X., and Boisvert, F. M. (2020) UBB pseudogene 4 encodes functional ubiquitin variants. Nature communications 11, 1306

26. Redman, K. L., and Rechsteiner, M. (1989) Identification of the long ubiquitin extension as ribosomal protein S27a. Nature 338, 438–440

27. Webb, G. C., Baker, R. T., Coggan, M., and Board, P. G. (1994) Localization of the human UBA52 ubiquitin fusion gene to chromosome band 19p13.1-p12. Genomics 19, 567–569

28. Kerscher, O., Felberbaum, R., and Hochstrasser, M. (2006) Modification of proteins by ubiquitin and ubiquitin-like proteins. Annual review of cell and developmental biology 22, 159–180

29. Komander, D., and Rape, M. (2012) The ubiquitin code. Annu Rev Biochem 81, 203–229

30. Hershko, A., and Ciechanover, A. (1998) The ubiquitin system. Annu Rev Biochem 67, 425–479

31. Ciechanover, A. (2005) Proteolysis: from the lysosome to ubiquitin and the proteasome. Nat Rev Mol Cell Biol 6, 79–87

32. Hershko, A., and Ciechanover, A. (1982) Mechanisms of intracellular protein breakdown. Annu Rev Biochem 51, 335–364

33. Bedford, L., Lowe, J., Dick, L. R., Mayer, R. J., and Brownell, J. E. (2011) Ubiquitin-like protein conjugation and the ubiquitin-proteasome system as drug targets. Nature reviews. Drug discovery 10, 29–46

34. Cappadocia, L., and Lima, C. D. (2018) Ubiquitin-like Protein Conjugation: Structures, Chemistry, and Mechanism. Chem Rev 118, 889–918

35. Hochstrasser, M. (2009) Origin and function of ubiquitin-like proteins. Nature 458, 422–429

36. Sisu, C., Pei, B., Leng, J., Frankish, A., Zhang, Y., Balasubramanian, S., Harte, R., Wang, D., Rutenberg-Schoenberg, M., Clark, W., Diekhans, M., Rozowsky, J., Hubbard, T., Harrow, J., and Gerstein, M. B. (2014) Comparative analysis of pseudogenes across three phyla. Proc Natl Acad Sci U S A 111, 13361–13366

37. Tonner, P., Srinivasasainagendra, V., Zhang, S., and Zhi, D. (2012) Detecting transcription of ribosomal protein pseudogenes in diverse human tissues from RNA-seq data. BMC genomics 13, 412

38. Wilson, D. N., and Doudna Cate, J. H. (2012) The structure and function of the eukaryotic ribosome. Cold Spring Harbor perspectives in biology 4

39. Tschochner, H., and Hurt, E. (2003) Pre-ribosomes on the road from the nucleolus to the cytoplasm. Trends Cell Biol 13, 255–263

40. Zemp, I., and Kutay, U. (2007) Nuclear export and cytoplasmic maturation of ribosomal subunits. Febs Lett 581, 2783–2793

41. Strunk, B. S., Loucks, C. R., Su, M., Vashisth, H., Cheng, S., Schilling, J., Brooks, C. L., 3rd, Karbstein, K., and Skiniotis, G. (2011) Ribosome assembly factors prevent premature translation initiation by 40S assembly intermediates. Science 333, 1449–1453

42. Ferreira-Cerca, S., Poll, G., Gleizes, P. E., Tschochner, H., and Milkereit, P. (2005) Roles of eukaryotic ribosomal proteins in maturation and transport of pre-18S rRNA and ribosome function. Mol Cell 20, 263–275

43. Cole, S. E., LaRiviere, F. J., Merrikh, C. N., and Moore, M. J. (2009) A convergence of rRNA and mRNA quality control pathways revealed by mechanistic analysis of nonfunctional rRNA decay. Mol Cell 34, 440–450

44. Strunk, B. S., Novak, M. N., Young, C. L., and Karbstein, K. (2012) A translation-like cycle is a quality control checkpoint for maturing 40S ribosome subunits. Cell 150, 111–121

45. Heyer, E. E., and Moore, M. J. (2016) Redefining the Translational Status of 80S Monosomes. Cell 164, 757–769

46. Biever, A., Glock, C., Tushev, G., Ciirdaeva, E., Dalmay, T., Langer, J. D., and Schuman, E. M. (2020) Monosomes actively translate synaptic mRNAs in neuronal processes. Science 367

47. Lopes, A. M., Miguel, R. N., Sargent, C. A., Ellis, P. J., Amorim, A., and Affara, N. A. (2010) The human RPS4 paralogue on Yq11.223 encodes a structurally conserved ribosomal protein and is preferentially expressed during spermatogenesis. BMC molecular biology 11, 33

48. Li, H., Huo, Y., He, X., Yao, L., Zhang, H., Cui, Y., Xiao, H., Xie, W., Zhang, D., Wang, Y., Zhang, S., Tu, H., Cheng, Y., Guo, Y., Cao, X., Zhu, Y., Jiang, T., Guo, X., Qin, Y., and Sha, J. (2022) A male germ-cell-specific ribosome controls male fertility. Nature 612, 725–731

49. Ghulam, M. M., Catala, M., and Abou Elela, S. (2020) Differential expression of duplicated ribosomal protein genes modifies ribosome composition in response to stress. Nucleic Acids Res 48, 1954–1968

50. Xue, S., and Barna, M. (2012) Specialized ribosomes: a new frontier in gene regulation and organismal biology. Nat Rev Mol Cell Biol 13, 355–369

51. Norris, K., Hopes, T., and Aspden, J. L. (2021) Ribosome heterogeneity and specialization in development. Wiley interdisciplinary reviews. RNA 12, e1644

52. Andres, O., Kellermann, T., Lopez-Giraldez, F., Rozas, J., Domingo-Roura, X., and Bosch, M. (2008) RPS4Y gene family evolution in primates. BMC evolutionary biology 8, 142

53. Chaillou, T. (2019) Ribosome specialization and its potential role in the control of protein translation and skeletal muscle size. Journal of applied physiology 127, 599–607

54. Karro, J. E., Yan, Y., Zheng, D., Zhang, Z., Carriero, N., Cayting, P., Harrrison, P., and Gerstein, M. (2007) Pseudogene.org: a comprehensive database and comparison platform for pseudogene annotation. Nucleic Acids Res 35, D55–60

55. Zhang, Z., Carriero, N., Zheng, D., Karro, J., Harrison, P. M., and Gerstein, M. (2006) PseudoPipe: an automated pseudogene identification pipeline. Bioinformatics 22, 1437–1439

56. Brunet, M. A., Brunelle, M., Lucier, J. F., Delcourt, V., Levesque, M., Grenier, F., Samandi, S., Leblanc, S., Aguilar, J. D., Dufour, P., Jacques, J. F., Fournier, I., Ouangraoua, A., Scott, M. S., Boisvert, F. M., and Roucou, X. (2019) OpenProt: a more comprehensive guide to explore eukaryotic coding potential and proteomes. Nucleic Acids Res 47, D403–D410

57. Michel, A. M., Fox, G., A, M. K., De Bo, C., O’Connor, P. B., Heaphy, S. M., Mullan, J. P., Donohue, C. A., Higgins, D. G., and Baranov, P. V. (2014) GWIPS-viz: development of a ribo-seq genome browser. Nucleic Acids Res 42, D859–864

58. Busch, H., and Goldknopf, I. L. (1981) Ubiquitin - protein conjugates. Molecular and cellular biochemistry 40, 173–187

59. Hershko, A., Ciechanover, A., and Rose, I. A. (1981) Identification of the active amino acid residue of the polypeptide of ATP-dependent protein breakdown. J Biol Chem 256, 1525–1528

60. Brunet, M. A., Lucier, J. F., Levesque, M., Leblanc, S., Jacques, J. F., Al-Saedi, H. R. H., Guilloy, N., Grenier, F., Avino, M., Fournier, I., Salzet, M., Ouangraoua, A., Scott, M. S., Boisvert, F. M., and Roucou, X. (2021) OpenProt 2021: deeper functional annotation of the coding potential of eukaryotic genomes. Nucleic Acids Res 49, D380–D388

61. Wen, B., Wang, X., and Zhang, B. (2019) PepQuery enables fast, accurate, and convenient proteomic validation of novel genomic alterations. Genome research 29, 485–493

62. Wen, B., Li, K., Zhang, Y., and Zhang, B. (2020) Cancer neoantigen prioritization through sensitive and reliable proteogenomics analysis. Nature communications 11, 1759

63. Rabl, J., Leibundgut, M., Ataide, S. F., Haag, A., and Ban, N. (2011) Crystal structure of the eukaryotic 40S ribosomal subunit in complex with initiation factor 1. Science 331, 730–736

64. Nicchitta, C. V., Lerner, R. S., Stephens, S. B., Dodd, R. D., and Pyhtila, B. (2005) Pathways for compartmentalizing protein synthesis in eukaryotic cells: the template-partitioning model. Biochemistry and cell biology = Biochimie et biologie cellulaire 83, 687–695

65. Stephens, S. B., and Nicchitta, C. V. (2008) Divergent regulation of protein synthesis in the cytosol and endoplasmic reticulum compartments of mammalian cells. Molecular biology of the cell 19, 623–632

66. Anger, A. M., Armache, J. P., Berninghausen, O., Habeck, M., Subklewe, M., Wilson, D. N., and Beckmann, R. (2013) Structures of the human and Drosophila 80S ribosome. Nature 497, 80–85

67. Spahn, C. M., Beckmann, R., Eswar, N., Penczek, P. A., Sali, A., Blobel, G., and Frank, J. (2001) Structure of the 80S ribosome from Saccharomyces cerevisiae--tRNA-ribosome and subunit-subunit interactions. Cell 107, 373–386

68. Montellese, C., van den Heuvel, J., Ashiono, C., Dorner, K., Melnik, A., Jonas, S., Zemp, I., Picotti, P., Gillet, L. C., and Kutay, U. (2020) USP16 counteracts mono-ubiquitination of RPS27a and promotes maturation of the 40S ribosomal subunit. eLife 9

69. Plassart, L., Shayan, R., Montellese, C., Rinaldi, D., Larburu, N., Pichereaux, C., Froment, C., Lebaron, S., O’Donohue, M. F., Kutay, U., Marcoux, J., Gleizes, P. E., and Plisson-Chastang, C. (2021) The final step of 40S ribosomal subunit maturation is controlled by a dual key lock. eLife 10

70. Sturm, M., Cheng, J., Bassler, J., Beckmann, R., and Hurt, E. (2017) Interdependent action of KH domain proteins Krr1 and Dim2 drive the 40S platform assembly. Nature communications 8, 2213

71. Hu, L., Wang, J., Liu, Y., Zhang, Y., Zhang, L., Kong, R., Zheng, Z., Du, X., and Ke, Y. (2011) A small ribosomal subunit (SSU) processome component, the human U3 protein 14A (hUTP14A) binds p53 and promotes p53 degradation. J Biol Chem 286, 3119–3128

72. Scott, W. A., Dhanji, E. Z., Dyakov, B. J. A., Dreseris, E. S., Asa, J. S., Grange, L. J., Mirceta, M., Pearson, C. E., Stewart, G. S., Gingras, A. C., and Campos, E. I. (2021) ATRX proximal protein associations boast roles beyond histone deposition. Plos Genet 17, e1009909

73. Clancy, A., Heride, C., Pinto-Fernandez, A., Elcocks, H., Kallinos, A., Kayser-Bricker, K. J., Wang, W., Smith, V., Davis, S., Fessler, S., McKinnon, C., Katz, M., Hammonds, T., Jones, N. P., O’Connell, J., Follows, B., Mischke, S., Caravella, J. A., Ioannidis, S., Dinsmore, C., Kim, S., Behrens, A., Komander, D., Kessler, B. M., Urbe, S., and Clague, M. J. (2021) The deubiquitylase USP9X controls ribosomal stalling. J Cell Biol 220

74. Warner, J. R., Knopf, P. M., and Rich, A. (1963) A multiple ribosomal structure in protein synthesis. Proc Natl Acad Sci U S A 49, 122–129

75. King, H. A., and Gerber, A. P. (2016) Translatome profiling: methods for genome-scale analysis of mRNA translation. Briefings in functional genomics 15, 22–31

76. Garshott, D. M., Sundaramoorthy, E., Leonard, M., and Bennett, E. J. (2020) Distinct regulatory ribosomal ubiquitylation events are reversible and hierarchically organized. eLife 9

77. 77. Slavov, N., Semrau, S., Airoldi, E., Budnik, B., and van Oudenaarden, A. (2015) Differential Stoichiometry among Core Ribosomal Proteins. Cell reports 13, 865–873

78. Yu, X., and Warner, J. R. (2001) Expression of a micro-protein. J Biol Chem 276, 33821–33825

79. Hong, S., Cho, Y. W., Yu, L. R., Yu, H., Veenstra, T. D., and Ge, K. (2007) Identification of JmjC domain-containing UTX and JMJD3 as histone H3 lysine 27 demethylases. Proc Natl Acad Sci U S A 104, 18439–18444

80. Xiang, Y., Zhu, Z., Han, G., Lin, H., Xu, L., and Chen, C. D. (2007) JMJD3 is a histone H3K27 demethylase. Cell Res 17, 850–857

81. De Santa, F., Totaro, M. G., Prosperini, E., Notarbartolo, S., Testa, G., and Natoli, G. (2007) The histone H3 lysine-27 demethylase Jmjd3 links inflammation to inhibition of polycomb-mediated gene silencing. Cell 130, 1083–1094

82. Agger, K., Cloos, P. A., Rudkjaer, L., Williams, K., Andersen, G., Christensen, J., and Helin, K. (2009) The H3K27me3 demethylase JMJD3 contributes to the activation of the INK4A-ARF locus in response to oncogene- and stress-induced senescence. Genes Dev 23, 1171–1176

83. Barradas, M., Anderton, E., Acosta, J. C., Li, S., Banito, A., Rodriguez-Niedenfuhr, M., Maertens, G., Banck, M., Zhou, M. M., Walsh, M. J., Peters, G., and Gil, J. (2009) Histone demethylase JMJD3 contributes to epigenetic control of INK4a/ARF by oncogenic RAS. Genes Dev 23, 1177–1182

84. Akiyama, T., Wakabayashi, S., Soma, A., Sato, S., Nakatake, Y., Oda, M., Murakami, M., Sakota, M., Chikazawa-Nohtomi, N., Ko, S. B., and Ko, M. S. (2016) Transient ectopic expression of the histone demethylase JMJD3 accelerates the differentiation of human pluripotent stem cells. Development 143, 3674–3685

85. Zhao, W., Li, Q., Ayers, S., Gu, Y., Shi, Z., Zhu, Q., Chen, Y., Wang, H. Y., and Wang, R. F. (2013) Jmjd3 inhibits reprogramming by upregulating expression of INK4a/Arf and targeting PHF20 for ubiquitination. Cell 152, 1037–1050

86. Sui, A., Xu, Y., Pan, B., Guo, T., Wu, J., Shen, Y., Yang, J., Guo, X. (2019) Histone demethylase KDM6B regulates 1,25-dihydroxyvitamin D3-induced senescence in glioma cells. Journal of cellular physiology 234, 17990–17998

87. Cao, R., Wang, L., Wang, H., Xia, L., Erdjument-Bromage, H., Tempst, P., Jones, R. S., and Zhang, Y. (2002) Role of histone H3 lysine 27 methylation in Polycomb-group silencing. Science 298, 1039–1043

88. Lavarone, E., Barbieri, C. M., and Pasini, D. (2019) Dissecting the role of H3K27 acetylation and methylation in PRC2 mediated control of cellular identity. Nature communications 10, 1679

89. Dellino, G. I., Schwartz, Y. B., Farkas, G., McCabe, D., Elgin, S. C., and Pirrotta, V. (2004) Polycomb silencing blocks transcription initiation. Mol Cell 13, 887–893

90. Bracken, A. P., Kleine-Kohlbrecher, D., Dietrich, N., Pasini, D., Gargiulo, G., Beekman, C., Theilgaard-Monch, K., Minucci, S., Porse, B. T., Marine, J. C., Hansen, K. H., and Helin, K. (2007) The Polycomb group proteins bind throughout the INK4A-ARF locus and are disassociated in senescent cells. Genes Dev 21, 525–530

91. Li, H., Zhang, H., Huang, G., Bing, Z., Xu, D., Liu, J., Luo, H., and An, X. (2022) Loss of RPS27a expression regulates the cell cycle, apoptosis, and proliferation via the RPL11-MDM2-p53 pathway in lung adenocarcinoma cells. Journal of experimental & clinical cancer research: CR 41, 33

92. Sun, X. X., DeVine, T., Challagundla, K. B., and Dai, M. S. (2011) Interplay between ribosomal protein S27a and MDM2 protein in p53 activation in response to ribosomal stress. J Biol Chem 286, 22730–22741

93. Becker, J., Barysch, S. V., Karaca, S., Dittner, C., Hsiao, H. H., Berriel Diaz, M., Herzig, S., Urlaub, H., and Melchior, F. (2013) Detecting endogenous SUMO targets in mammalian cells and tissues. Nat Struct Mol Biol 20, 525–531

94. Bouchard, D., Wang, W., Yang, W. C., He, S., Garcia, A., and Matunis, M. J. (2021) SUMO paralogue-specific functions revealed through systematic analysis of human knockout cell lines and gene expression data. Molecular biology of the cell 32, 1849–1866

95. Samir, P., Browne, C. M., Rahul, Sun, M., Shen, B., Li, W., Frank, J., and Link, A. J. (2018) Identification of Changing Ribosome Protein Compositions using Mass Spectrometry. Proteomics 18, e1800217

96. Serrano, M., Hannon, G. J., and Beach, D. (1993) A new regulatory motif in cell-cycle control causing specific inhibition of cyclin D/CDK4. Nature 366, 704–707

97. Stock, J. K., Giadrossi, S., Casanova, M., Brookes, E., Vidal, M., Koseki, H., Brockdorff, N., Fisher, A. G., and Pombo, A. (2007) Ring1-mediated ubiquitination of H2A restrains poised RNA polymerase II at bivalent genes in mouse ES cells. Nat Cell Biol 9, 1428–1435

98. Jacobs, J. J., Kieboom, K., Marino, S., DePinho, R. A., and van Lohuizen, M. (1999) The oncogene and Polycomb-group gene bmi-1 regulates cell proliferation and senescence through the ink4a locus. Nature 397, 164–168

99. Frankish, A., Diekhans, M., Ferreira, A. M., Johnson, R., Jungreis, I., Loveland, J., Mudge, J. M., Sisu, C., Wright, J., Armstrong, J., Barnes, I., Berry, A., Bignell, A., Carbonell Sala, S., Chrast, J., Cunningham, F., Di Domenico, T., Donaldson, S., Fiddes, I. T., Garcia Giron, C., Gonzalez, J. M., Grego, T., Hardy, M., Hourlier, T., Hunt, T., Izuogu, O. G., Lagarde, J., Martin, F. J., Martinez, L., Mohanan, S., Muir, P., Navarro, F. C. P., Parker, A., Pei, B., Pozo, F., Ruffier, M., Schmitt, B. M., Stapleton, E., Suner, M. M., Sycheva, I., Uszczynska-Ratajczak, B., Xu, J., Yates, A., Zerbino, D., Zhang, Y., Aken, B., Choudhary, J. S., Gerstein, M., Guigo, R., Hubbard, T. J. P., Kellis, M., Paten, B., Reymond, A., Tress, M. L., and Flicek, P. (2019) GENCODE reference annotation for the human and mouse genomes. Nucleic Acids Res 47, D766–D773

100. Lam, H. Y., Khurana, E., Fang, G., Cayting, P., Carriero, N., Cheung, K. H., and Gerstein, M. B. (2009) Pseudofam: the pseudogene families database. Nucleic Acids Res 37, D738–743

101. Sayers, E. W., Beck, J., Brister, J. R., Bolton, E. E., Canese, K., Comeau, D. C., Funk, K., Ketter, A., Kim, S., Kimchi, A., Kitts, P. A., Kuznetsov, A., Lathrop, S., Lu, Z., McGarvey, K., Madden, T. L., Murphy, T. D., O’Leary, N., Phan, L., Schneider, V. A., Thibaud-Nissen, F., Trawick, B. W., Pruitt, K. D., and Ostell, J. (2020) Database resources of the National Center for Biotechnology Information. Nucleic Acids Res 48, D9–D16

102. Cunningham, F., Allen, J. E., Allen, J., Alvarez-Jarreta, J., Amode, M. R., Armean, I. M., Austine-Orimoloye, O., Azov, A. G., Barnes, I., Bennett, R., Berry, A., Bhai, J., Bignell, A., Billis, K., Boddu, S., Brooks, L., Charkhchi, M., Cummins, C., Da Rin Fioretto, L., Davidson, C., Dodiya, K., Donaldson, S., El Houdaigui, B., El Naboulsi, T., Fatima, R., Giron, C. G., Genez, T., Martinez, J. G., Guijarro-Clarke, C., Gymer, A., Hardy, M., Hollis, Z., Hourlier, T., Hunt, T., Juettemann, T., Kaikala, V., Kay, M., Lavidas, I., Le, T., Lemos, D., Marugan, J. C., Mohanan, S., Mushtaq, A., Naven, M., Ogeh, D. N., Parker, A., Parton, A., Perry, M., Pilizota, I., Prosovetskaia, I., Sakthivel, M. P., Salam, A. I. A., Schmitt, B. M., Schuilenburg, H., Sheppard, D., Perez-Silva, J. G., Stark, W., Steed, E., Sutinen, K., Sukumaran, R., Sumathipala, D., Suner, M. M., Szpak, M., Thormann, A., Tricomi, F. F., Urbina-Gomez, D., Veidenberg, A., Walsh, T. A., Walts, B., Willhoft, N., Winterbottom, A., Wass, E., Chakiachvili, M., Flint, B., Frankish, A., Giorgetti, S., Haggerty, L., Hunt, S. E., Gr, I. I., Loveland, J. E., Martin, F. J., Moore, B., Mudge, J. M., Muffato, M., Perry, E., Ruffier, M., Tate, J., Thybert, D., Trevanion, S. J., Dyer, S., Harrison, P. W., Howe, K. L., Yates, A. D., Zerbino, D. R., and Flicek, P. (2022) Ensembl 2022. Nucleic Acids Res 50, D988–D995

103. Wilkins, M. R., Gasteiger, E., Bairoch, A., Sanchez, J. C., Williams, K. L., Appel, R. D., and Hochstrasser, D. F. (1999) Protein identification and analysis tools in the ExPASy server. Methods in molecular biology 112, 531–552

104. Michel, A. M., Kiniry, S. J., O’Connor, P. B. F., Mullan, J. P., and Baranov, P. V. (2018) GWIPS-viz: 2018 update. Nucleic Acids Res 46, D823–D830

105. Brunet, M. A., and Roucou, X. (2019) Mass Spectrometry-Based Proteomics Analyses Using the OpenProt Database to Unveil Novel Proteins Translated from Non-Canonical Open Reading Frames. Journal of visualized experiments: JoVE

106. Cox, J., and Mann, M. (2008) MaxQuant enables high peptide identification rates, individualized p.p.b.-range mass accuracies and proteome-wide protein quantification. Nature biotechnology 26, 1367–1372

107. Tyanova, S., Temu, T., and Cox, J. (2016) The MaxQuant computational platform for mass spectrometry-based shotgun proteomics. Nature protocols 11, 2301–2319

108. Wieczorek, S., Combes, F., Lazar, C., Giai Gianetto, Q., Gatto, L., Dorffer, A., Hesse, M., Coute, Y., Ferro, M., Bruley, C., and Burger, T. (2017) DAPAR & ProStaR: software to perform statistical analyses in quantitative discovery proteomics. Bioinformatics 33, 135–136

109. Giai Gianetto, Q., Combes, F., Ramus, C., Bruley, C., Coute, Y., and Burger, T. (2016) Calibration plot for proteomics: A graphical tool to visually check the assumptions underlying FDR control in quantitative experiments. Proteomics 16, 29–32

110. Huez, I., Creancier, L., Audigier, S., Gensac, M. C., Prats, A. C., and Prats, H. (1998) Two independent internal ribosome entry sites are involved in translation initiation of vascular endothelial growth factor mRNA. Mol Cell Biol 18, 6178–6190

111. Martineau, Y., Le Bec, C., Monbrun, L., Allo, V., Chiu, I. M., Danos, O., Moine, H., Prats, H., and Prats, A. C. (2004) Internal ribosome entry site structural motifs conserved among mammalian fibroblast growth factor 1 alternatively spliced mRNAs. Mol Cell Biol 24, 7622–7635

112. Erales, J., Marchand, V., Panthu, B., Gillot, S., Belin, S., Ghayad, S. E., Garcia, M., Laforets, F., Marcel, V., Baudin-Baillieu, A., Bertin, P., Coute, Y., Adrait, A., Meyer, M., Therizols, G., Yusupov, M., Namy, O., Ohlmann, T., Motorin, Y., Catez, F., and Diaz, J. J. (2017) Evidence for rRNA 2’-O-methylation plasticity: Control of intrinsic translational capabilities of human ribosomes. Proc Natl Acad Sci U S A 114, 12934–12939

113. Chen, S., Zhou, Y., Chen, Y., and Gu, J. (2018) fastp: an ultra-fast all-in-one FASTQ preprocessor. Bioinformatics 34, i884–i890

114. Trapnell, C., Williams, B. A., Pertea, G., Mortazavi, A., Kwan, G., van Baren, M. J., Salzberg, S. L., Wold, B. J., and Pachter, L. (2010) Transcript assembly and quantification by RNA-Seq reveals unannotated transcripts and isoform switching during cell differentiation. Nature biotechnology 28, 511–515

115. Bray, N. L., Pimentel, H., Melsted, P., and Pachter, L. (2016) Near-optimal probabilistic RNA-seq quantification. Nature biotechnology 34, 525–527

116. Soneson, C., Love, M. I., and Robinson, M. D. (2015) Differential analyses for RNA-seq: transcript-level estimates improve gene-level inferences. F1000Research 4, 1521

117. Love, M. I., Huber, W., and Anders, S. (2014) Moderated estimation of fold change and dispersion for RNA-seq data with DESeq2. Genome biology 15, 550

118. Chang, S., Park, B., Choi, K., Moon, Y., Lee, H. Y., and Park, H. (2016) Hypoxic reprograming of H3K27me3 and H3K4me3 at the INK4A locus. Febs Lett 590, 3407–3415

119. Hellemans, J., Mortier, G., De Paepe, A., Speleman, F., and Vandesompele, J. (2007) qBase relative quantification framework and software for management and automated analysis of real-time quantitative PCR data. Genome biology 8, R19

120. Brosseau, J. P., Lucier, J. F., Lapointe, E., Durand, M., Gendron, D., Gervais-Bird, J., Tremblay, K., Perreault, J. P., and Elela, S. A. (2010) High-throughput quantification of splicing isoforms. Rna 16, 442–449

121. Petryszak, R., Keays, M., Tang, Y. A., Fonseca, N. A., Barrera, E., Burdett, T., Fullgrabe, A., Fuentes, A. M., Jupp, S., Koskinen, S., Mannion, O., Huerta, L., Megy, K., Snow, C., Williams, E., Barzine, M., Hastings, E., Weisser, H., Wright, J., Jaiswal, P., Huber, W., Choudhary, J., Parkinson, H. E., and Brazma, A. (2016) Expression Atlas update--an integrated database of gene and protein expression in humans, animals and plants. Nucleic Acids Res 44, D746–752

